# Biologically plausible learning in a deep recurrent spiking network

**DOI:** 10.1101/613471

**Authors:** David Rotermund, Klaus R. Pawelzik

**Affiliations:** University of Bremen, Insitute for Theoretical Physics, Bremen, Germany

## Abstract

Artificial deep convolutional networks (DCNs) meanwhile beat even human performance in challenging tasks. Recently DCNs were shown to also predict real neuronal responses. Their relevance for understanding the neuronal networks in the brain, however, remains questionable. In contrast to the unidirectional architecture of DCNs neurons in cortex are recurrently connected and exchange signals by short pulses, the action potentials. Furthermore, learning in the brain is based on local synaptic mechanisms, in stark contrast to the global optimization methods used in technical deep networks. What is missing is a similarly powerful approach with spiking neurons that employs local synaptic learning mechanisms for optimizing global network performance. Here, we present a framework consisting of mutually coupled local circuits of spiking neurons. The dynamics of the circuits is derived from first principles to optimally encode their respective inputs. From the same global objective function a local learning rule is derived that corresponds to spike-timing dependent plasticity of the excitatory inter-circuit synapses. For deep networks built from these circuits self-organization is based on the ensemble of inputs while for supervised learning the desired outputs are applied in parallel as additional inputs to output layers.

Generality of the approach is shown with Boolean functions and its functionality is demonstrated with an image classification task, where networks of spiking neurons approach the performance of their artificial cousins. Since the local circuits operate independently and in parallel, the novel framework not only meets a fundamental property of the brain but also allows for the construction of special hardware. We expect that this will in future enable investigations of very large network architectures far beyond current DCNs, including also large scale models of cortex where areas consisting of many local circuits form a complex cyclic network.

## Introduction

Since the pioneering work of McCullogh and Pitts [1] a range of paradigmatic models were introduced for understanding neuronal computations in the brain. Despite unrealistic architectures and simplified neuron models feed-forward networks helped to understand receptive fields in early areas of visual cortex and recurrent networks prepared the ground for understanding universal properties of memory. With stochastic nodes they served as role model for probabilistic computations [2]. For modeling realistic architectures hybrid models as e.g. restricted Boltzmann Machines and Helmholtz Machines [3] combine hierarchical architectures with recurrent interactions. While theoretical neuroscience fleshed out these formal models by demonstrating that their core properties are conserved in models with realistic neurons and synapses, artificial neuronal networks deviated more and more from the biological role model. In particular, the so called Deep Convolutional Networks (DCNs) that now form the core of many state of the art artificial intelligence systems [4–19] are quite far from biological reality. Nevertheless, DCNs were recently picked up by neuroscience [20] as a possible explanation of neuronal responses to natural stimuli in visual cortex [21–24].

DCNs lack important constraints of real neuronal networks. In particular, cortical networks have no simple hierarchy, neurons in the brain are recurrently connected and communicate with action potentials that are brief electrical pulses also called ‘spikes’. Also, learning in the brain is mostly realized by local synaptic mechanisms that rely only on pre- and postsynaptic activity, as e.g. in spike-timing dependent plasticity (STDP) [25–27]. This stands in stark contrast to the usual algorithms for learning in DCNs. In particular, the back-propagation learning algorithm used there requires specific error signals to bridge many layers in acyclic network architectures ([5, 28], https://www.tensorflow.org). Last not least, it is not obvious which functions neuronal networks in the brain actually serve. Many networks in the brain might underly even several different computational functions, depending on the tasks at hand, context, and state of attention.

There have been many attempts to bridge the gap between artificial neuronal networks and realistic models of the brain. For instance back-propagation was related to realistic conditions [29–32] and recurrent networks were proposed for learning [3, 33–37]. These models, however, are still rather abstract, with states that are usually taken to be also the signals exchanged among neurons, with unrealistic constraints on the weights and unrealistic learning rules [33, 38].

Brains consist of interconnected areas. On a finer level each area is composed of recurrently interacting local circuits. It is believed that cortex is composed of microcircuits that have a rather stereotyped architecture with thousands mutually coupled neurons [39]. It is not known how the efficacies of the excitatory synapses connecting areas and microcircuits are self-organizing in a way that promotes function and ensures dynamical stability.

A promising approach for understanding cortical microcircuits are the so called generative models [40–42] that can be realized by neuronal networks performing sparse coding. Sparse coding neuronal circuits (SCNC) perform compressed sampling, a highly efficient and mathematically well understood data compression principle [43] which e.g. successfully reproduced simple cell responses in visual cortex [44, 45].

It was repeatedly shown that networks with spiking neurons can realize SCNCs [43, 46–58]. It is, however, not clear how to combine sparse coding circuits to build large recurrent networks with many layers that have the potential to explain neuronal responses and have competitive computational performance.

Here, we present a general framework for realistic networks of sparse coding neuronal circuits that takes relevant constraints into account including recurrent architectures, signaling by noisy spikes, locality of synaptic learning mechanisms and Dale’s law for long range excitatory couplings. It has repeatedly been demonstrated that SCNCs can be realized by biologically realistic models of spiking neurons [46, 47]. The current approach skips detailed modeling of the SCNCs and instead jumps to an abstract formulation introduced in [58] by which the inference is iteratively performed with each spike received by a population of neurons termed ‘inference population’ (IP). Here, for convenience, we briefly review the dynamics of latent variables and the learning of weights in this model from [58]:

A IP consists of a population of *N*_*g*_ neurons which represents its input pattern *µ* by finding an estimate of the probability *P*_*g′*,*µ*_(*s*_*g′*_) for receiving spikes from neurons *s*_*g′*_ originating from another population *g′* of *N*_*g′*_ input neurons. This estimate 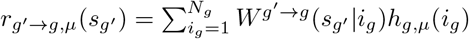 is represented by a latent variable *h*_*g,µ*_(*i*_*g*_). The weights *W*^*g′→g*^(*s*_*g′*_|*i*_*g*_) as well as the latent variables *h*_*g,µ*_(*i*_*g*_) are positive numbers from the range [0, 1]. Furthermore, we enforce normalization of *h*_*g,µ*_(*i*_*g*_) over *i*_*g*_ (i.e. the population’s neurons) and of *W*^*g′→g*^(*s*_*g′*_|*i*_*g*_) over the input neurons *s*_*g′*_. In [58] it was shown that this approach is consistent if the input neurons fire independently with a Poisson point process. Then *r*_*g′→g,µ*_(*s*_*g′*_) is an estimate of the probability *P*_*g′*,*µ*_(*s*_*g′*_) of receiving the next spike from input node *s*_*g′*_.

For a single IP, the dynamics of the latent variables *h*_*g,µ*_(*i*_*g*_) as well as learning rules for the weights *W*^*g→gI*^(*s*_*g′*_|*i*_*g*_) were derived in [58] from maximizing the negative log likelihood which is equivalent to maximizing the cross-entropy *E*_*g′→g*_ between *P*_*g′,µ*_(*s*_*g′*_) and *r*_*g′→g,µ*_(*s*_*g′*_) over all *M* patterns

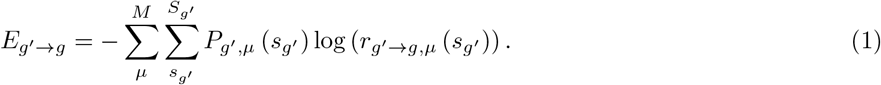

The dynamics of the latent variables *h*_*g,µ*_(*i*_*g*_) for optimizing *E*_*g′→g*_ in expectation with each input spike 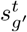 reads

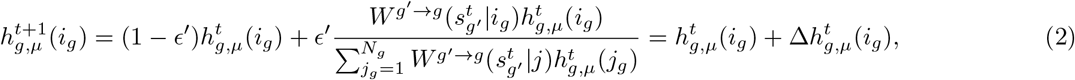

where *ϵ′* parametrizes a low-pass filter. This multiplicative spike-by-spike dynamics respects positivity and normalization. It can be considered a formal realization of the computations actually occurring in real neuronal circuits that perform inference for which several biologically more realistic implementations are available [46, 47]. From maximizing the cross-entropy, it is also possible to derive learning rules that are online or batch learning rules and can be multiplicative or additive. The multiplicative online-learning rule conserves positivity and updates the weights *W*^*g′→g*^(*s*|*i*) with every spike 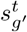 according to

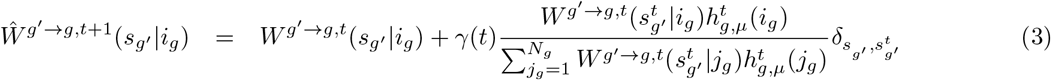

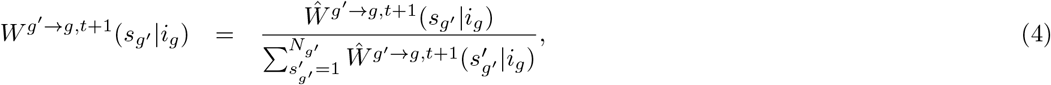

with *γ*(*t*) as a learning rate. Note that in this formulation *t* represents the iteration steps that are incremented with each spike event encountered by the IP thereby replacing real time. Therefore its dynamics is termed ‘Spike-by-Spike’ update.

Rearrangement by using the spike-by-spike dynamics of the latent variables to express the weight changes reveals that weight changes can be expressed by

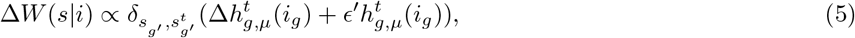

and subsequent normalization that keeps the weights bounded. In other words, the learning rule combines contributions from a differential Hebbian rule and a Hebbian term. Since differential Hebbian learning with spikes is related to spike-timing dependent plasticity [59], the spike based learning of the synaptic weights providing the inputs into a IP can be considered a variant of spike-timing dependent plasticity [25–27].

After developing the general framework for networks of these IPs, we show that they can realize Boolean functions which demonstrates the generality of the approach. Using the architecture of deep convolutional networks and MNIST handwritten digits benchmark (http://yann.lecun.com/exdb/mnist/), we then show that it achieves performances that approach those of their technical cousins, if they are used with similar learning techniques and network structure. Finally, we use the same network architecture as used for handwritten digit classification network to mimic a visual cortex model by unsupervised training on natural images. We find that it reproduces realistic receptive fields RFs and also some context effects observed in area V1.

## Results

### Harmony dynamics

We here introduce arbitrary networks of *G* inference populations (Fig. 1). As in [58] each IP consists of *N*_*g*_ neurons, where *g* = 1*, …, G* denotes the different populations. Also, each neuron in every IP carries a latent variable *h*_*g,µ*_(*i*_*g*_) that will also depend on (iteration) time. The index *µ* represents one input pattern from an ensemble with *M* patterns (i.e. *µ* = 1*, …, M*). The latent variables have non-negative values *h*_*g,µ*_(*i*_*g*_) ∈ [0, 1]. Also following [58], all latent variables within a IP are in a local competition via normalization 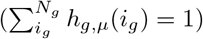. In the original Spike-by-Spike framework, spikes are simply the indices identifying which neuron *s*_*g′*_ in the input population *g′* is actually sending the spike. This happens at iteration step *t* which enumerates the emission events of spikes. Also here, as in the original SbS network, time can be progressed from one spike to the next emitted from the unique input population thereby replacing real time. Using this notation a IP receives input from other populations via spikes 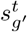. Note that in between spikes (i.e. in the real time interval from iteration steps *t* to *t* + 1), none of the entities in a SbS network will change.

**Fig 1.**
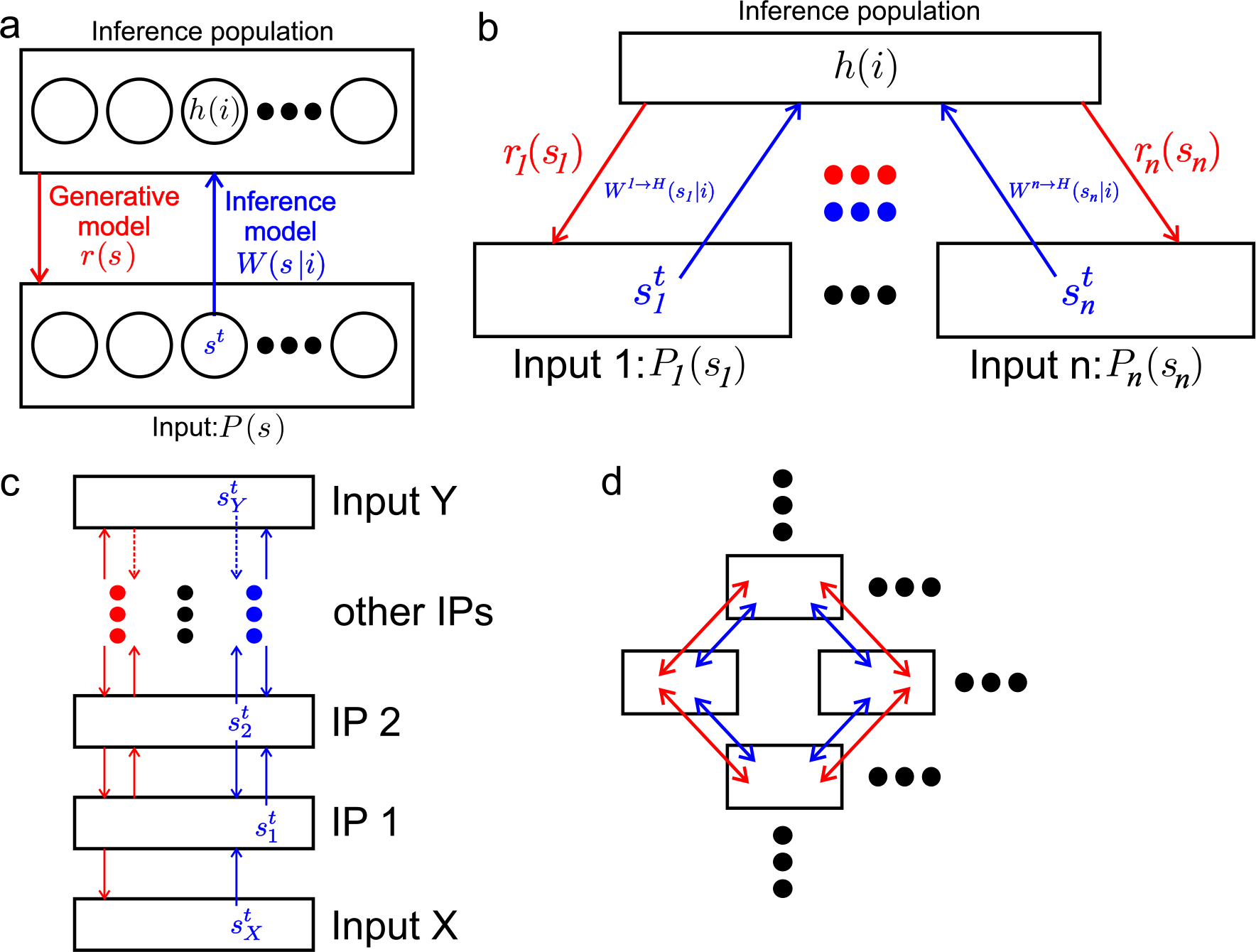
Constructing networks of inference populations (NIPs). a) The basic element of the proposed networks is the inference population (top box). With only one input population (lower box), the ensemble of *N* neurons in the IP develop a representation *r*(*s*) = Σ_*j*_ *W*(*s*|*j*)*h*(*j*) of the underlying input spike probabilities *P*(*s*) by iteratively updating their latent variables 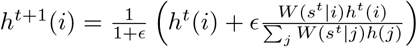 from each spike *s*^*t*^ received from the input population. This dynamics was derived from the gradient of the cross entropy *E* which quantifies the quality of the representation [58]. Since the IP develops a prediction of its inputs, it can be said to ‘explain’ it. b) When several input populations are providing input to an IP, the set of latent variables *h*(*i*) in the IP will seek a compromise where the spike probabilities in each of the *n* input populations are well explained. Assuming independence of the spike events in each input population, the cross entropy decomposes into separate summands for each input population. Thereby the contributions from each spike to the update for the latent variables will simply add up (Eq. 11). c) and d): When several IPs are mutually coupled to build networks, they not only perform inference on their respective inputs, but also generate stochastic spikes according to the values of their latent variables *h*(*i*). The probability for generating the respective next spike 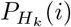 is set to *h*(*i*). Thereby a given IP sends information about its current state to the other IPs to which it is coupled. Since each IP seeks a consistent representation of its inputs, mutually coupled IPs will seek to ‘explain’ each other as well as the spikes from input populations if they are also coupled to the latter. c) A hierachical architecture with only one IP at each layer contains recurrent interactions among layers. d) More complex networks can contain motivs with loops of mutually explaining IPs.

For extending this idea to more than one IP providing input spikes to a different IP *t* can be used to enumerate all spikes in the network, independent from which IP they come. This allows for each population having different activity levels.

Each spike in each population becomes generated from the normalized rate *P*_*g′,µ*_(*s*_*g′*_) which has *N*_*g′*_ neurons itself and will in general depend also on time. For IPs within the network *g′*, *P*_*g′,µ*_(*s*_*g′*_) is given by *h*_*g′,µ*_(*s*_*g′*_) while for input layers *g*^∗^ we choose *I*_*g∗*_,_*µ*_(*s*_*g∗*_) to denote the stationary probability for the next spike emitted when a particular input pattern *µ* is present. For determining the next spike in the whole system one first has to determine which population will fire next depending on the activity levels *λ*_*g*_, and then stochastically select which neuron in this population will send the spike. That is, we have a doubly stochastic process which in real time would correspond to independent Poisson point processes.

Every IP attempts to represent the probability *P*_*g′*,*µ*_(*s*_*g′*_) to receive input spikes from a population *g′* by

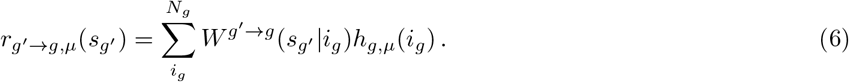

The receiving populations *g* collect information about *P*_*g′,µ*_(*s*_*g′*_) only through incoming spikes 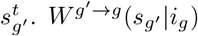 are weights that describe the connection strength between the neuron *s*_*g′*_ which emits the spike and neuron *i*_*g*_ that receives the spike. The interactions between IPs are considered to represent long range connections where the weights are non-negative numbers with *W* (*s*_*g′*_|*i*_*g*_) ∈ [0, 1]. In addition, the weights are normalized such that, if a connection between population *g′* and *g* exits at all, 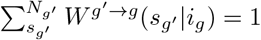.

In the following we derive an update rule for the latent variables in a IP *g* that allows to process the incoming spikes from all populations *g′* connected to *g* in an iterative fashion. We call this update ‘harmony dynamics’ since the latent variables in the receiving IP intuitively attempt to harmonize all the inputs, which it sees from the different input populations, simultaneously into one consistent explanation using the same set of latent variables.

Formally, we capture this intuition by the pairwise cross-entropy *E*_*g′→g*_ between *P*_*g′*_(*s*_*g′*_) and *r*_*g′→g,µ*_(*s*_*g′*_) over all *M* patterns by

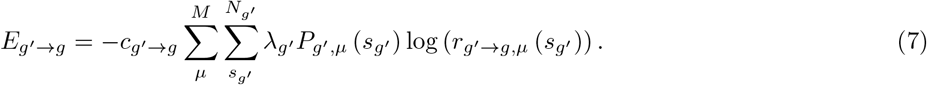

If, due to the selection of the network structure, there is no connection between population *g′* and population *g*, then *c*_*g′→g*_ is set to zero. Otherwise *c*_*g′→g*_ is set to one.

Assuming statistical independence of the spike events across populations, the total cross-entropy for the whole network with *G* inference populations is

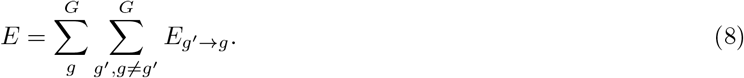

Based on *E*, we can now calculate the gradient for *h*_*g,µ*_(*u*_*g*_)

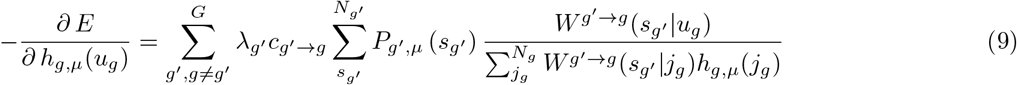

and for *W*^*g′→g*^(*u*_*g′*_|*v*_*g*_)

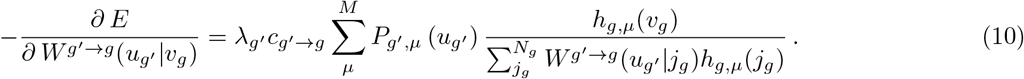

Both gradients can be used in a multiplicative gradient optimization algorithm:

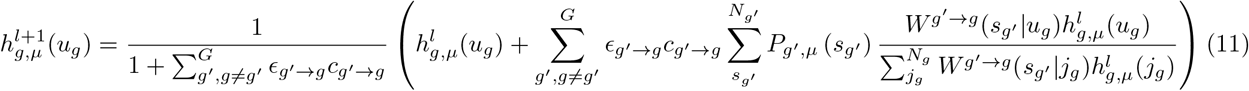

with *ϵ*_*g′→g*_ as a non-negative update rate. We will later use this parameter to absorb the activity levels of the respective populations.

For the weights we obtain

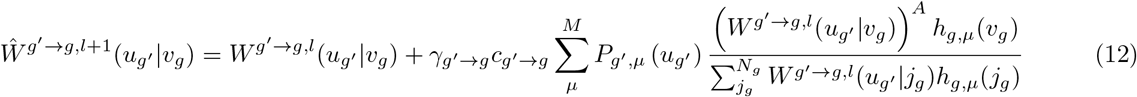

with *γ*_*g′→g*_ as non-negative learning rate that can also be used to absorb the activity levels of the populations. *A* = 1 results in a multiplicative learning rule and *A* = 0 is for an additive learning rule. To keep the normalization of the weights intact, an additional normalization step is required:

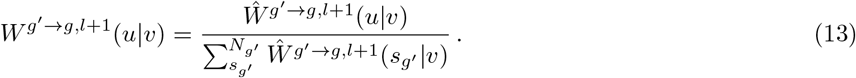

For an iterative update of the latent variables with every single incoming spike, only the single spike 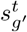 from *P*_*g′,µ*_(*s*_*g′*_) is used (which corresponds to 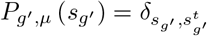 and 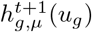 is updated with one spike at a time. Thus equation 11 can be simplified into

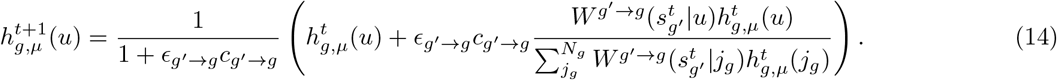

While equation 12 represents a batch learning rule which operates on many spikes and patterns, an online learning rule can easily be derived with 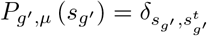:

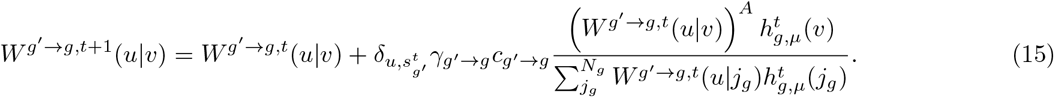

For keeping the weights normalized, equation 13 needs to be applied subsequently too. It should to be noted that *P*_*g′,µ*_(*s*_*g′*_) = *h*_*g′,µ*_(*s*_*g′*_) if the spike originates from a inference population or is *P*_*g′,µ*_(*s*_*g′*_) = *I*_*g′,µ*_(*s*_*g′*_) if the spikes are generated by neurons without latent variables but with a probability distribution *I*_*g′,µ*_(*s*_*g′*_), like it would be the case in an input layer.

For making these equations more vivid, let us assume a network with four layers: One input-layer *X* with an input probability distribution *p*_*X*_ (*s*_*X*_), two hidden layers *H*1 with latent variables *h*_*H*1_(*i*_*H*1_) and *H*2 with latent variables *h*_*H*2_(*i*_*H*2_) and an output layer *HY* with latent variables *h*_*HY*_ (*i*_*HY*_). Layers *H*1, *H*2, and *HY* are one IP each. Furthermore, the network structure is set such that spikes 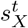 from input layer *X* are sent to *H*1 and also layer *H*2 sends spikes 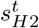 to layer *H*1. Layer *H*2 receives spikes 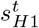 from layer *H*1 as well as spikes 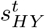 from the output layer. And finally the output layer *HY* collects spikes 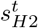 from layer *H*2.

If a spike 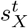 is generated by *p*_*X*_(*s*_*X*_) then *h*_*H*1,*µ*_(*u*_*H*1_) is updated as follows:

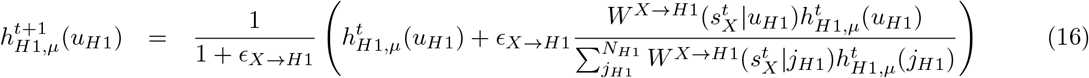

A spike 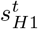 produced by *h*_*H*1,*µ*_(*u*_*H*1_) elicits an update in *h*_*H*2,*µ*_(*u*_*H*2_):

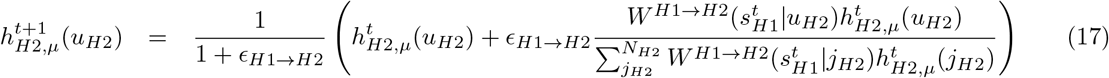

The two layers *H*1 and *HY* need to be updated if *H*2 produces a spike 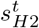 from 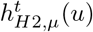:

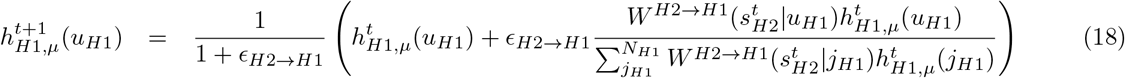

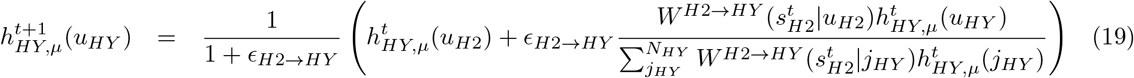

And finally, if output layer *HY* produces a spikes 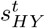 from *h*_*HY,µ*_(*u*_*HY*_) then *h*_*H*2,*µ*_(*u*_*H*2_) needs an update via

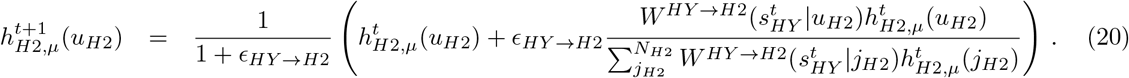

Concerning the weights, the batch learning rules are simplified into

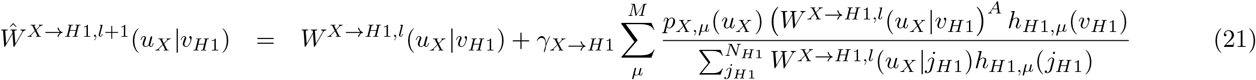

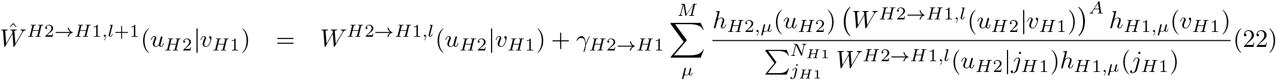

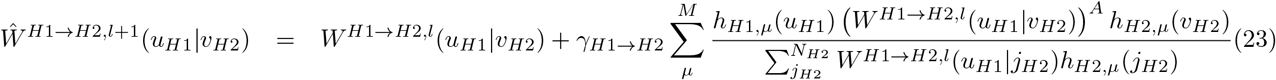

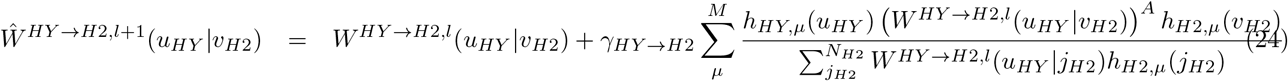

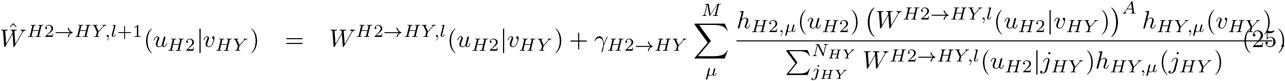

The corresponding weights 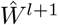 are re-normalized after every update. It should be noted that in some instances *p*_*X,µ*_(*u*_*X*_), *h*_*H*1,*µ*_(*u*_*H*1_), *h*_*H*2*,µ*_(*u*_*H*2_), and *h*_*HY,µ*_(*u*_*HY*_) can be replaced by an estimated probability distribution from the observed spikes (e.g. 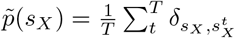 if *T* spikes from the input layer *X* have been observed.)

### SbS network for calculating parity functions

#### Simulations with perfect weights

Generality of the framework follows from its ability to realize and learn arbitrary Boolean functions in the spirit of McCullogh&Pitts [1]. In [58] we already showed that flat one layer SbS networks can learn any Boolean function, however, very inefficiently. Here we demonstrate this for far more efficient hierarchical architectures using the example of parity functions which essentially count the number of input bits with value 1. If the sum is even, these functions result in 0 otherwise they return 1. For two bits this corresponds to the XOR function. For these type of networks, every input bit as well as the output bit are encoded by a pair of neurons each. During processing of a given pattern, the firing probabilities of the input neurons are fixed. The activity of the first neuron is for the logical value 0, while the activity of the second neuron represents the logical state 1. For each pair of input neurons, the activity is mutually exclusive because an input bit can only be 0 or 1. While the desired output would also show the same mutual exclusivity, in simulations the outcome can be less perfect. Thus selecting the identity of the output neuron with the higher activity (with regard to its *h*(*i*)-values or their spike counts) is used for decode the output bit. Figure 2(a) shows the hierarchical network constructed for the XOR task. On the lowest layer, two input bits are shown. The boxes signal that all the neurons in that box are in a IP, where the sum over their h-values, and thus the neuron’s firing probabilities, is one. Two of these populations generate the input spikes for this network.

**Fig 2.**
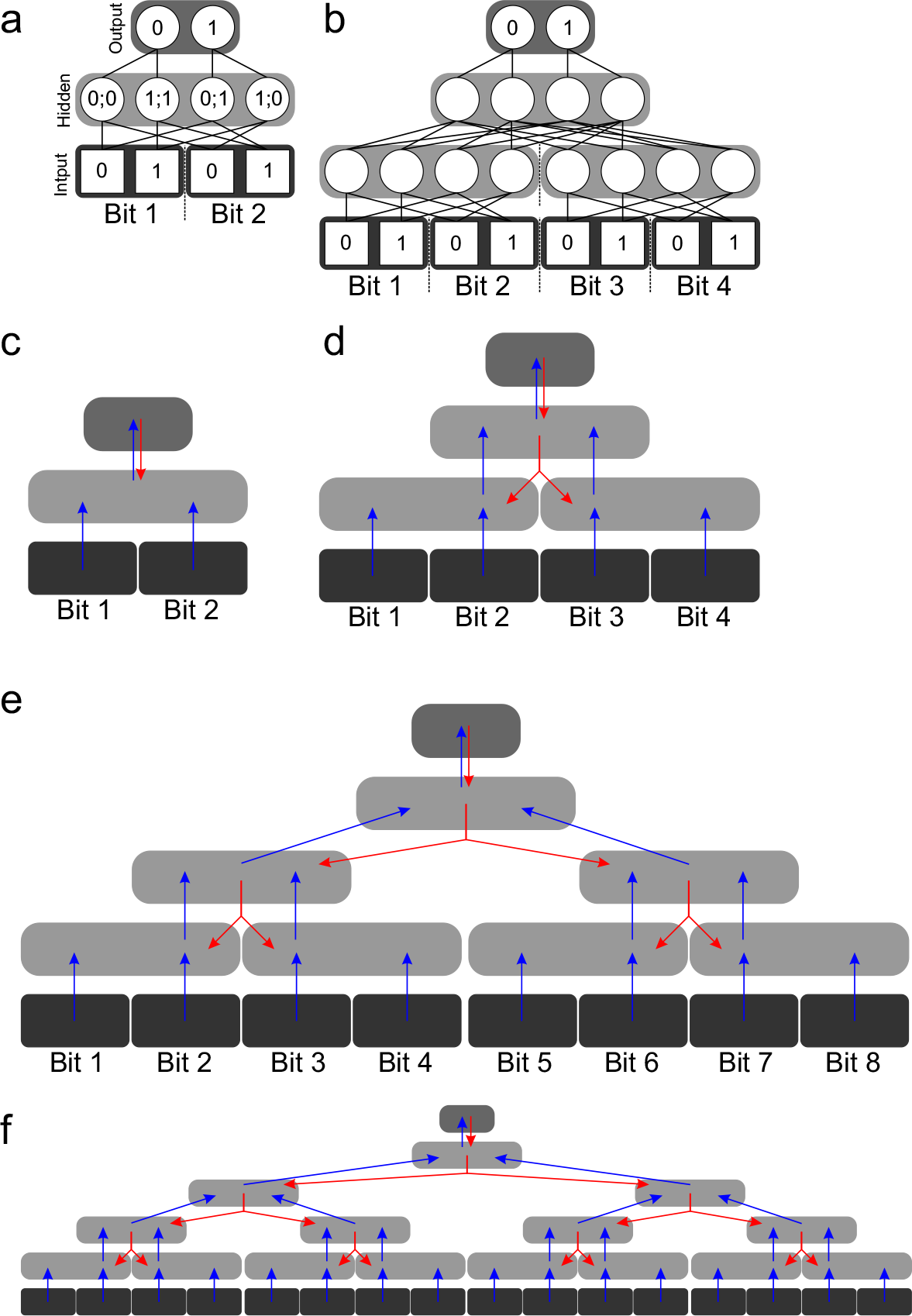
Structures of the parity networks. A: Structure of the network for the XOR-problem. B: Network with two hidden layer for solving the 4 bit parity task. C-F: The flow of information via spikes between the different IPs for the XOR problem as well as the 4, 8, and 16 bit parity problem. Blue arrows represent spikes that transfer information further away from the input layer, while red arrows transport information in the other direction. Split arrows represent spikes that are evaluated by two IPs.

The hidden layer, which consists of four neurons, processes these incoming input spikes accordingly the SbS h-dynamic. Since this dynamic realizes a winner-take-all behavior, the h-values will end in a very sparse distribution. For every one of the four possible input bit combinations, only one hidden neuron will be active while the other three neurons will be silent. The bit combination that will activate a hidden neuron is noted inside its circle. Furthermore the hidden layer will get additional spikes from the output layer, which are also incorporated into the h-values of the hidden neurons. The arrows in figure 2(c) shows where the information flows, via the spikes, through out the XOR network.

In figure 2, the mathematically correct and ‘active’ (*W* (*s|i*) *>* 0) weights are represented by lines while weights with value of 0 (’inactive’) are not shown. These weights are used for both directions of information flow. The values of active weights bridging two layers have always the same value. These values depend, due to the normalization of weights Σ_*s*_*W*(*s|i*) = 1, on the direction of information flow and the corresponding two layers. E.g. the shown weights from the hidden layer to the output neurons have to a value of one, while for the other direction the weights have a value of 0.5.

The network for the 4 bit parity-problem figure2(b) was constructed by copying the input and the hidden layer of the XOR network twice and placed side-by-side. Now the input layer represents 4 bits via four IPs. Both IPs in the first hidden layer individually react to their input bit patterns as shown in the hidden layer of figure 2(a). A second hidden layer is added for combining the spike activities of the two parts of the lower hidden layer. Again the second hidden layer shows a winner-take-all dynamic. Table 1 shows the input bit combinations to which the neurons of the second hidden layer react to. On top of this second hidden layer a output layer, similar to the XOR version, is placed.

**Table 1.**
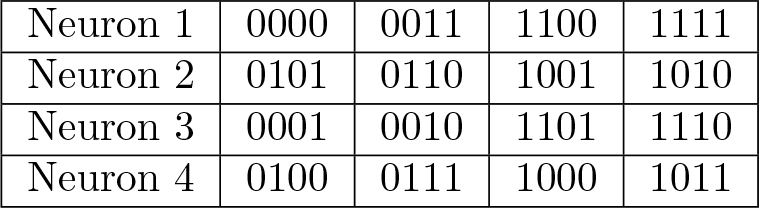
List of input bit patterns that the neurons in the second hidden layer of the 4 bit parity problem react to.

Instead generating only a single spike in the whole network, we for the following simulations let every IP produce one spike each at each iteration step, for reasons of computational efficiency. In fact, this procedure achieves equivalent results if the rates *ϵ* for updating the latent variables are chosen according to the respective activity levels which in the present case a taken to be equal. While the flow of the spikes in the XOR network figure 2(c) is very simple, the 4 bit parity network it is more complex. The information flow, shown in figure 2(d), can be read as follows: Blue arrows represent spikes flowing forward while red spikes depict spikes progressing backwards. Red split arrows are processed by two IPs at the same time. The red and blue arrow leaving one IP represent the same spike.

The 4 bit parity function can be easily extended to a 8 bit (figure 2(e)) or a 16 bit (figure 2(e)) parity function. Doubling the number of input neurons adds an extra hidden layer to the parity network. A 32 bit network has 5 hidden layers, a 64 bit network has 6 hidden layers, and a 128 bit has 7 hidden layers.

Figure 3 reports the performance of these networks using the correct weights. As measure for the performance, always the *h*-value of the output neuron that is representing the wrong logic output is analyzed. The desired *h*-value response for this neuron is 0. Shown is the distance in dependency in the number of spikes each IP has produced, while *h* starts as uniform distribution (which is a value of 0.5 for the output layer). Figure 3 shows that the deeper the networks get, the more spikes are required for a correct response. Furthermore, the simulations reveal that deeper networks require a smaller *ϵ*, otherwise the higher layer would get already stuck in sparse *h*-distributions before the information from the input layer can reach them. Due to the multiplicative nature of the *h*-dynamic, the algorithm has problems to recover from states where the *h*-value of significant neurons for the pattern meet the value of zero. A smaller *ϵ* slows this process down until the input information has propagated to where it is needed. The simulations show that with enough spikes these networks deliver the correct response for all the tested parity functions.

**Fig 3.**
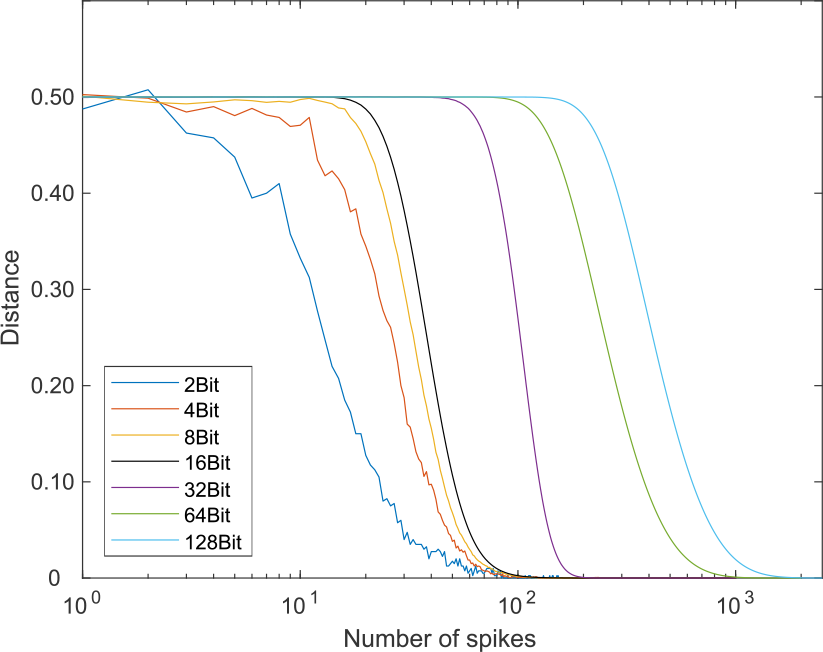
Performance of parity networks with different bit widths. The average distance for the neuron from the output layer that represents the wrong output logic state is shown. The distance is calculated between the desired output *h* value for this neuron (which is 0) and the observed *h* value during simulating these networks. The distance is shown in dependence of the number of spikes generated by each IP. The simulations start with uniform distribution for the *h*-values for all the IPs. All shown curves are averages over 100 realizations with different random seeds for the simulations. For the networks with bit widths of 2, 4, 8, and 16 *ϵ* = 0.1 is used. For larger parity functions, smaller values for *ϵ* are required (32 bit: *ϵ* = 0.03, 64 bit: *ϵ* = 0.02, and 128 bit: *ϵ* = 0.012). The same *ϵ* value is used for the forward and backward transport of information. The deeper the network, the more spikes are required to propagate the information through the network. For the networks with 32 input bits and more, only a random subset of patterns is tested (32 bit: 2^16^ patterns, 64 bit: 2^15^ patterns, and 128 bit: 2^14^ patterns). This is necessary since the required computing time was too high.

#### Robustness to noise on the weights

We checked the robustness to additive positive noise on the weights for the XOR case (Figure 4). For noise levels *<* 1, the weights values of the correct weights which were originally zero (’inactive’ weight values) are guaranteed to be still smaller than the weight values which were non-zero (’active’ weight values) in the correct weights. For noise levels *>* 1, former ‘inactive’ weight values can become larger than the ‘active’ weight values. Until a noise level of 0.4 for the XOR network and 0.27 for the four bit parity network, the networks can still perform their task perfectly (tested after 1000 spikes per IP for every pattern and *ϵ* = 0.1). With higher noise levels, the performance gradually degrades.

**Fig 4.**
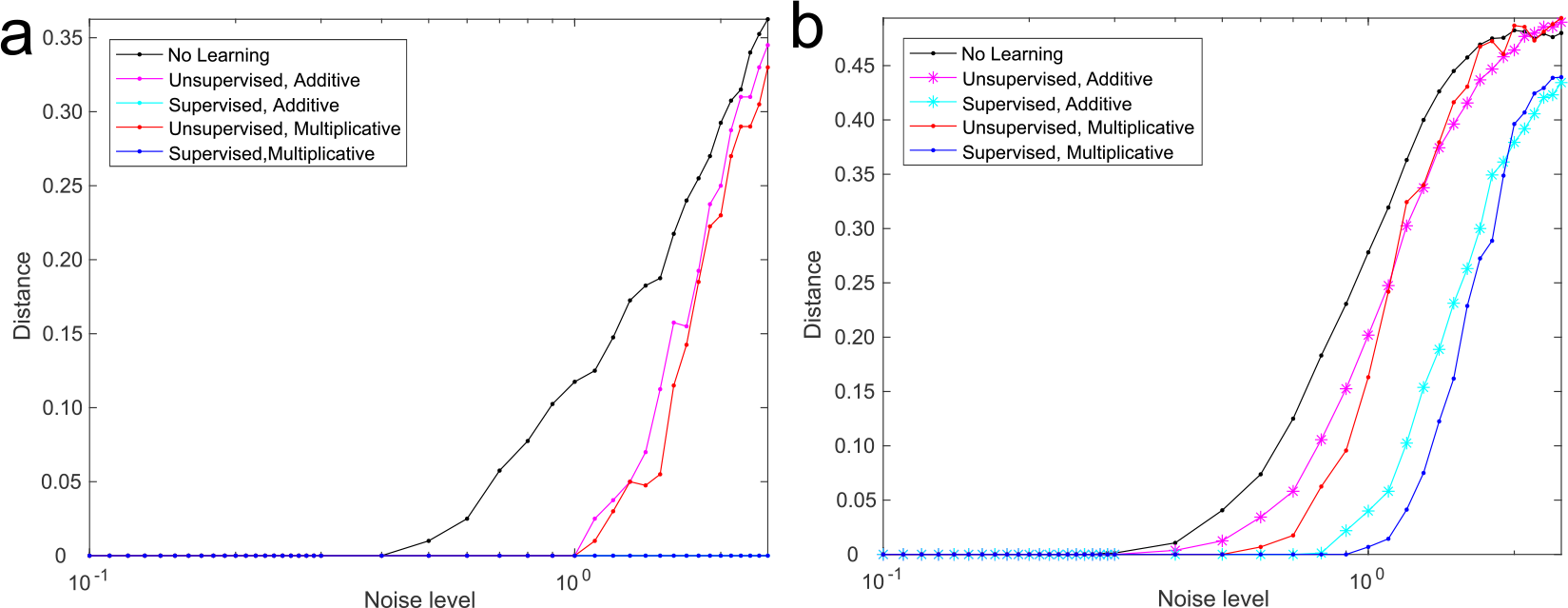
Performance under noise on the weights. While figure 3 uses mathematically perfect weights, these two sub-figures (a: XOR, b: 4 bit parity) explore the decay of the performance under noise on the weights. The weights are perturbed with additive positive uniform noise. At a noise level of 1, the amplitude of the noise and the correct ‘active’ weight values can have the same amplitude (’active’ refers to the weight values for the correct weights that are larger zero and ‘inactive’ corresponds to weights values of zeros for the correct weights). Below a noise level of 1, all the disturbed ‘active’ weights are still guaranteed to be lager than the ‘inactive’ weights. Above a noise level of 1, ‘inactive’ weights can get larger than the ‘active’ weights. The black lines show that the network can handle a moderate amount of noise on its own as well as that the performance starts to decay under higher noise levels. The red curves show the performances after an unsupervised learning. In these cases no information about the correct output was given while both online-learning rules (magenta: additive and dark red: multiplicative) tried to repair the error. This leads to a moderate reduction in error. The blue lines show the performance when the same procedure is performed under supervised learning. In these conditions, information about the correct output was given during learning. In the case of the XOR network, this procedure can repair the weights perfectly. The cyan (additive online-learning) and dark blue (multiplicative online-learning) are both lying on the x-axes. For the four bit parity network, the performance enhancement is less pronounced. This is due to local minima. All the shown performances values are an average over 100 initial conditions.

We also tested if unsupervised learning (without information about the correct output) can counter-act otherwise destructive perturbations of the weights. For this purpose we apply on-line learning, where with every spike *s*^*t*^ the weights are updated according to

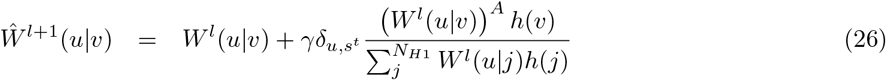

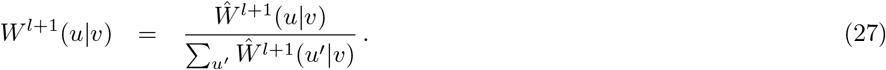

In the simulations (*ϵ* = 0.1) the first 900 spikes were used to allow the network to leave the initial state. Then the weights are updated with every spike for the next 100 spikes. This is done for all the patterns and this procedure is repeated for 100 times. The curves in figure 4 show this repair attempt (*γ*_*Add*_ = 0.00001, *γ*_*Multi*_ = 0.0005). In the case of the XOR function, the network is able to repair itself for noise-levels of up to 1. This is the noise level where ‘inactive’ weight values can get larger than ‘active’ values. This approach is less successful for the 4 bit parity function. The additive on-line learning rule (*γ*_*Add*_ = 0.00001) can repair the network until a noise level of 0.3. The multiplicative on-line learning rule (*γ*_*Multi*_ = 0.0001) can keep up until a noise level of 0.5.

Figure 4 also shows what happens if supervised on-line learning is used. The procedure is identical to the unsupervised on-line learning with the difference that during learning the *h*-distribution of the output layer is fixed to the correct output distribution. Now the additive (*γ*_*Add*_ = 0.005) and the multiplicative learning rule (*γ*_*Multi*_ = 0.005) can restore the performance of the XOR network for the whole range of tested noise levels (tested up to a noise level of 2.5). For the four bit function, this translates into a noise level of 0.7 for the additive learning rule (*γ*_*Add*_ = 0.00001) and 0.9 for the multiplicative learning rule (*γ*_*Multi*_ = 0.001).

#### Learning weights from scratch

After using perfect or by noise disturbed weights, the question remains if it is possible to learn the weights from scratch. Starting with uniform weights with, an additional positive uniformly distributed noise of 1%, the presented supervised on-line strategies were applied. For the XOR network, using the multiplicative learning rule with *γ*_*Multi*_ = 0.005, a perfect classification performance was reached after four learning steps for all 100 initial random initializations. The classification performance measures if the *h*-value of the output neuron representing the correct result has a high *h*-value than the other output neuron for the wrong answer.

For the four bit parity function learning is not that easy. For example with the multiplicative learning rule we weren’t able to find *γ* values that allowed to learn all 100 initial conditions perfectly. Figure 5 shows the performance learning the weights with the additive on-line learning rule for this four bit network. We found that it is necessary to change the learning rate *γ* over the learning steps. If learning was started with a small *γ*, the network state seem to get stuck in local minima. If *γ* was big, then escaping those local minima was possible but the learning rule wasn’t able to refine the weights such that a good performance was reached after learning. Thus we applied a strategy which used a big *γ* in the beginning and reduced its values over the learning steps. Figure 5 shows that it is possible to learn the weights perfectly (tested with 100 initial conditions) with changing *γ* gradually or step-wise. Beside changing the *γ* values over time and starting with random weights, the procedures described for figure 4 were used.

**Fig 5.**
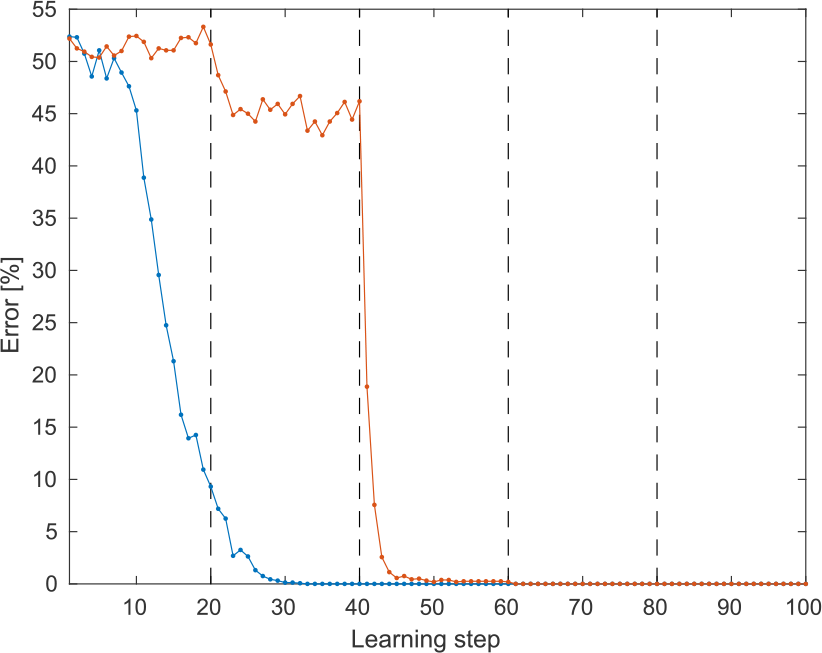
Online-learning weights for the 4 bit parity function from scratch. After initialization of the the weight (uniform values with a 1% noise carpet for breaking symmetries), the additive online learning rule is applied in a supervised fashion. Figure 4 showed that local minima present a challenge during learning. Thus it is necessary to either reduce the learning parameter *γ* smoothly (blue curve; *γ* = 0.01 *•* 5^*−l/*20^ with *l* as the learning step number, starting with 1 and *N*_*Spikes*_ = 500) or shown in the red curve, slowly increasing the number of spikes (*N*_*Spikes*_ = 100 + *l ⋅* 10) while simultaneously jumping from higher learning rates *γ* to smaller values (changes of *γ* occur at the dashed lines; *γ*_0_ = 0.01, *γ*_20_ = 0.005, *γ*_40_ = 0.001, *γ*_60_ = 0.0005 and *γ*_80_ = 0.0001). The shown performance values are averaged over 100 initial conditions. The results for multiplicative online-learning are not shown because the network wasn’t always able to find the perfect weights for the all 100 initial conditions. The learning curves for the XOR case are not shown because after just four learning steps the correct weights had been found.

On-line learning is, due to the normalization steps after every spike which was used for updating the weights, highly computational demanding. Instead we wanted to test if batch learning, using the information from all patterns for one weight update, can be used as an alternative. One variant for a batch learning rule is

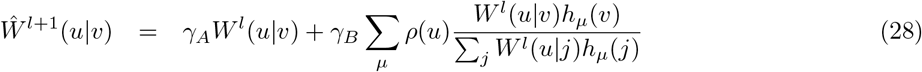

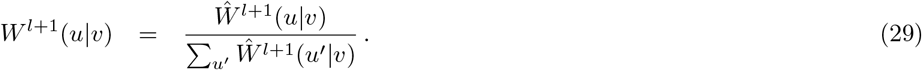

*ρ*(*u*) is the *h*-value distribution from the respective spike emitting population to *h*_*µ*_(*v*). Figure 6a shows the result for this learning rule (*γ*_*A*_ = 0.1 and *γ*_*B*_ = 1). However, learning was difficult. This time, the problem with local minima were even stronger. Starting with the simulation procedure from figure 5 and applying the batch learning rule, several changes were necessary: *ϵ* was changed with every one spikes of the 500 spikes (per IP) according *ϵ* = 0.1 *⋅ t*^−1/3^ with *t* as the number of processed spikes. *h*_*µ*_(*v*) and *ρ*(*u*) used in the learning rule were averaged over the last 50 spikes. During learning, the *h*-values of the output layer weren’t fixed as before and was progressing according the normal *h*-dynamics. For representing the correct class, for every spike the output layer sends to its neighboring layer, an additional spike from the correct output neuron was injected and processed. Learned are the weights from the input layer to the first hidden layer, the weights from the first hidden layer to the second hidden layer, and the weights from the output layer to the second hidden layer. The other weights are transposed and re-normalized versions of the their corresponding weights in the other direction. Even though all these modification were done, the problem with the local minima largely remained. Some of the 100 initial conditions ended up in the correct weights but not all. Thus a schema was developed that resets part of the networks weights back to random values and continues learning with them. Beginning with fully random weights, after every 25 learning steps it is tested if this network is able to do the desired task perfectly. If it isn’t able to do so, the weights between the first and the second hidden layer are randomly reset. After the network is still not functional after 50 learning steps, all weights except the ones between the input and the first hidden layers are reset to random values. Then, after an additional 25 learning steps, all weights of an unsuccessful learning attempt were randomly reset and the cycle began anew. Figure 6a shows that after several of these cycles all 100 initial conditions reached correct weights. It is important to note: In a more general task, typically the test if training was successful would be done on the training data. However, here the training data and the test data is, due to the structure of the used task, identically. This can be seen as an unfair mixing of training and test data but can’t be avoided in this example.

**Fig 6.**
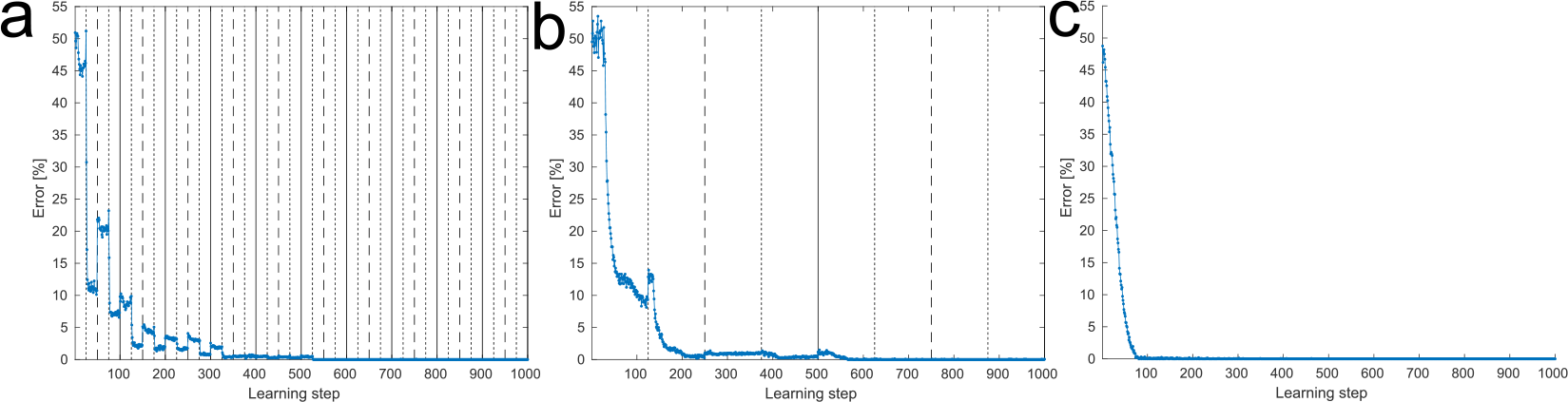
Batch-learning weights for the 4 bit parity function from scratch. Like in figure 5, learning started with randomly initialized weights. Three approaches of batch learning are shown. a: Normal batch learning using *ϵ* = 0.1 ⋅ *t*^−1/3^ with *t* as the amount of spikes produced by every individual IP. For ensuring that all of the 100 tested initial conditions, over which the shown performance is averaged, end in a perfect set of weights, it was necessary to introduce a reset schema to the reanimate networks that were trapped in local minima. After every learning step a performance test was done. In the case were the output wasn’t perfect, part or all weights of the network were reset and learning continued. At all types of black lines, the weights between the two hidden layer were reset. At the dashed and solid black lines, the weights between the output and its neighboring hidden layer are reset. And at the solid lines all weights are reset. b: Instead of the normal batch learning, a learning rule derivative from ADAM was used. Less resets are required. c: The normal batch learning rule is used but in combination with unsupervised pre-learning of the weights between input and first hidden layer as well as an offset in the spike generation is present. The latter one can be understood as a variant of annealing and ensures that the weights are slowly settling into their correct position. This offset keeps all neurons (to some degree) active and is linearly reduced over the learning steps until it reaches zero. Overall these examples show that batch learning is very sensitive to the initial conditions of the weights and effort is required to ensure that learning with all initial weight is successful.

Inspired by the Adam learning rule for deep networks [60], we modified the batch learning rule for figure 6b. Keeping the rest of the latter simulation the same, except extending the described reset periods from 25 to 125, we get ∆*W*^*l*^(*u|v*) as gradient after presenting every input pattern once for learning step *l*:

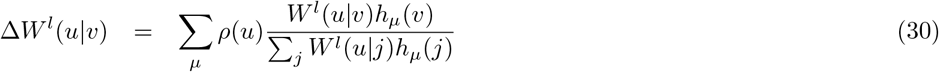

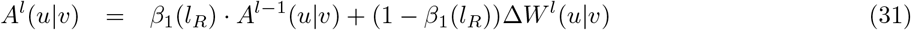

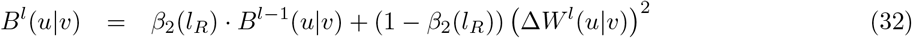

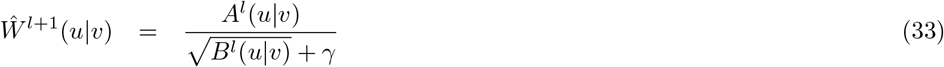

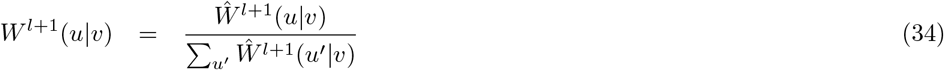

with

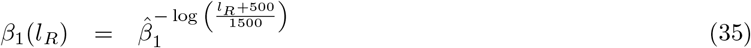

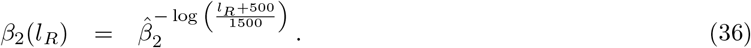

In the shown simulation the parameters were chosen as follows: *γ* = 10^−8^, 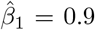, and 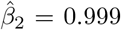. *l*_*R*_ is measured from the last random weight reset for that specific weight. Figure 6b shows that the number of resets is reduced and in the end all 100 initial random conditions are learned perfectly.

In figure 6c we tested how the learning performance is effected by unsupervised pre-learning, for the weights between the input layer and the first hidden layer, as well as ‘annealing’ the spike generation process. During unsupervised pre-learning, the network is reduced into its input layer and the first hidden layer. For 500 learning steps and 100 spikes per learning step and pattern, the learning rule presented in equation 28 is used (*γ*_*A*_ = 1 and *γ*_*B*_ = 0.1 while the last 100 spikes are used for averaging *ρ* and *h*). Then these pre-learned weights are copied into the full network. Now a learning procedure similar to the one described for figure 6a is applied (*γ*_*A*_ = 1 and *γ*_*B*_ = 0.45 with 2000 spikes per pattern. *h* is averaged over the last 200 spikes and *ρ*(*u*) is calculated by counting the number of spikes generated by the corresponding input neuron *u* and dividing it by the respective total spike count). However, in this case we applied a variant of annealing for the spike production process. Instead of using the corresponding *h* values or the input & output probability distribution directly, they got an offset which is depended of number of already performed learning steps *l* using the following equation:

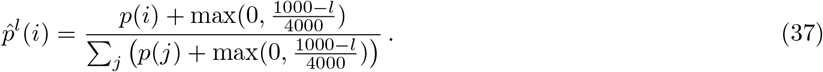

With this equation *p*(*i*) (which is *h*(*i*) for IPs or the input probability distribution for input populations) is transformed into 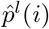, which is now used for drawing spikes. Using pre-learning and the presented annealing process allows to remove the prior used reset schema and producing correctly working weights for all 100 initial conditions. Furthermore, the learning rule is doing this faster than the other two approaches (see figure 6).

#### Filling in missing data

In the presented networks, information flows not only from the input to the output but also backwards from the output to the input. Thereby, the framework not only performs formal inference but can be used for associative replenishment of information missing in the input. For investigating this information flow in more detail, the 8 bit parity function network is used (see figure 7a). Given the output as well as seven of the input bits, the eighth input bit is defined by the 8 bit parity look-up table. For the simulations shown in figure 7, we added a new readout IP with two neurons that is observing the spikes emitted from the first hidden layer, using the transposed and re-normalized weights that the corresponding input bits would have. Figure 7b shows for two sets of *ϵ* parameters (*ϵ*_*Forward*_ = *ϵ*_*Backward*_ = 0.05 as well as *ϵ*_*Forward*_ = 0.1 and *ϵ*_*Backward*_ = 0.05) how the information about the missing bit is accumulated in the readout IP (the shown curves are averaged over 100 initial conditions), in dependency of the number of spikes processed by the readout population per pattern.

**Fig 7.**
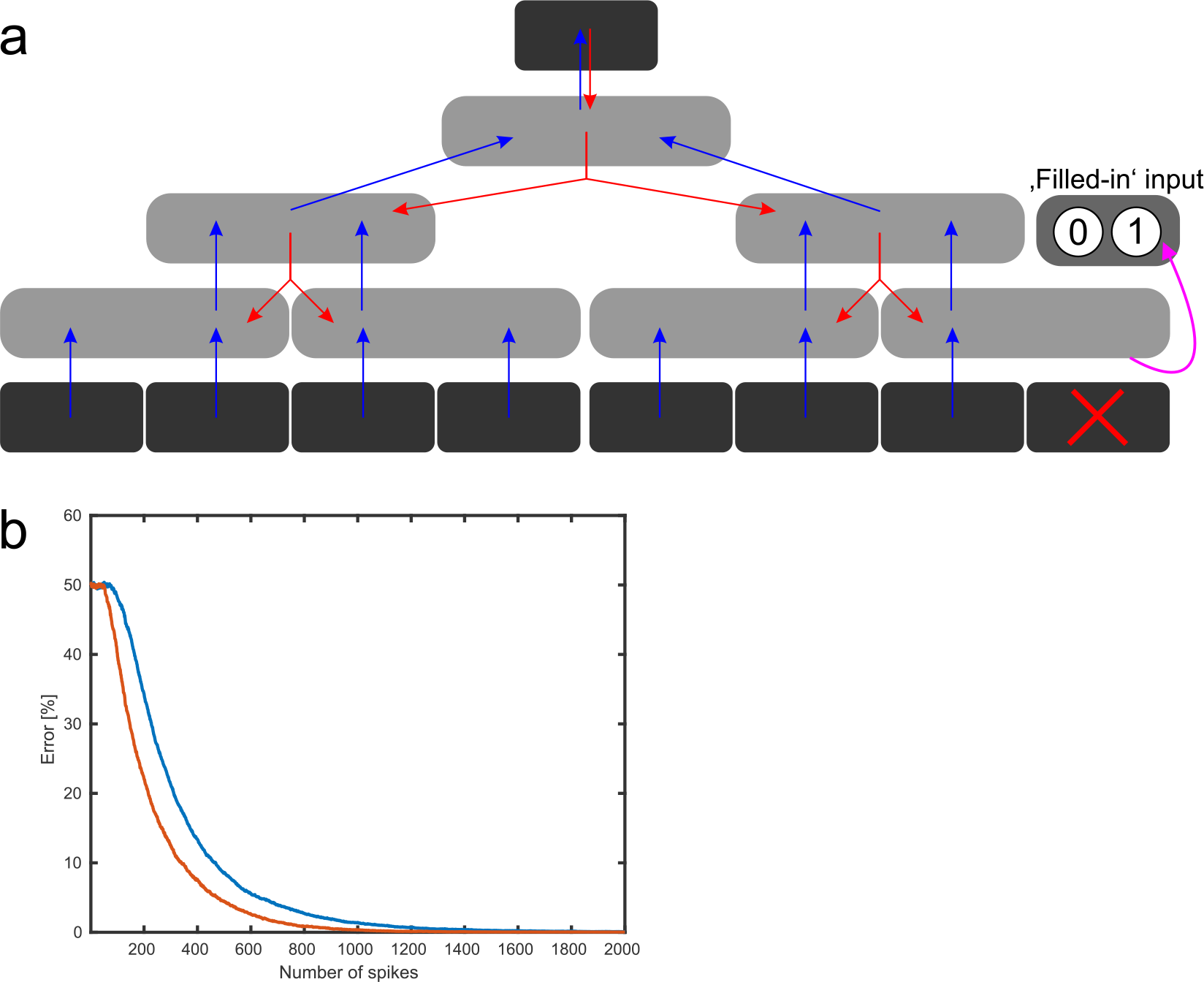
Filling in of missing input. a: From the 8 bit parity function one input bit is removed and the corresponding output value is applied to the output layer. This output, through the definition of the 8 bit parity function, selects a correct logical state for the missing input neuron. An additional normalization module with two neurons is added to the network, using the transposed and re-normalized weights to the removed input bit. This module observes the spikes from its hidden layer and uses its h-dynamic to infer which logical state for the missing input bit the network ‘imagines’. b: Quality of the filled input is shown over the number of processed spikes in the extra normalization module. The 7 bits of the input in combination with the output as context information, brings the network into a state of activity which allows a perfect reconstruction of the missing input (performance averaged over 100 initial conditions). For the blue cure *ϵ*_*Forward*_ = *ϵ*_*Backward*_ = 0.05 were used and for calculating the red curve *ϵ*_*Forward*_ was changed to 0.1.

#### Deep SbS networks – the MNIST example

The MNIST database (see http://yann.lecun.com/exdb/mnist/) is a commonly used toy example for machine learning (see Google’s Tensor Flow tutorial for MNIST: https://www.tensorflow.org/tutorials/estimators/cnn). The data set contains handwritten digits with 28 × 28 pixels with 8 bit gray values. It provides 60,000 training pattern and 10,000 test pattern. We took the network structure for a deep convolutional neuronal network (CNN) presented in the Tensor Flow tutorial and modified it according to our needs. The used network structure is shown in figure 8. There a three main differences in the network structure compared to the Tensor Flow tutorial: a.) The convolution layers in the Tensor Flow tutorial used padding with zeros. This keeps the input and output dimensions of convolution layers constant. In the case of the SbS network, we don’t propagate the information if part of the convolutional kernel is outside of input. The convolutional layers use 5 × 5 kernel. Thus we reduce the 28 × 28 pixel per input image to 24 × 24 pixel for the output of the first convolutional layer *H*1 and it creates a reduction from 12 × 12 pixel to 8 × 8 pixels for the second convolutional layer *H*3. b.) The pooling layers *H*2 and *H*4 are only using the competition between neurons in a IP inherent in the SbS model and no special max functions like in the Tensor Flow tutorial. This is realized by the following weight structure for *H*1 *→ H*2 and *H*3 *→ H*4: 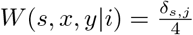 (with *x ∈* [1, 2] and *y ∈* [1, 2]). Using such a weight matrix lumps together inputs from different spatial coordinates but only from the same feature. However, it creates a competition between the different features. c.) The input pixel undergo a so-called ‘on/off’ split [58]. In practice the 8 bit pixel values *I*(*x, y*) are brought on a global *±*1 value range, where the center point of the scale is a 127.5 pixel value with 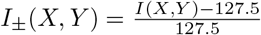. After that the split is done by *I*_*On*_(*x, y*) = [*I*_±_(*X, Y*)]_+_ and *I*_*Off*_(*x, y*) = [*−I_±_*(*X, Y*)]_+_ where [*…*]_+_ sets all negative values to zeros and keeps all positive values as they are. Furthermore, we don’t apply input distortion methods to increase the size of the training set by the input modifications. Like with the non zero padding approach for the convolution layer, this is a time saving strategy due to the much higher computational demand for simulating this type of neurons compared with a standard Tensor Flow MNIST network.

**Fig 8.**
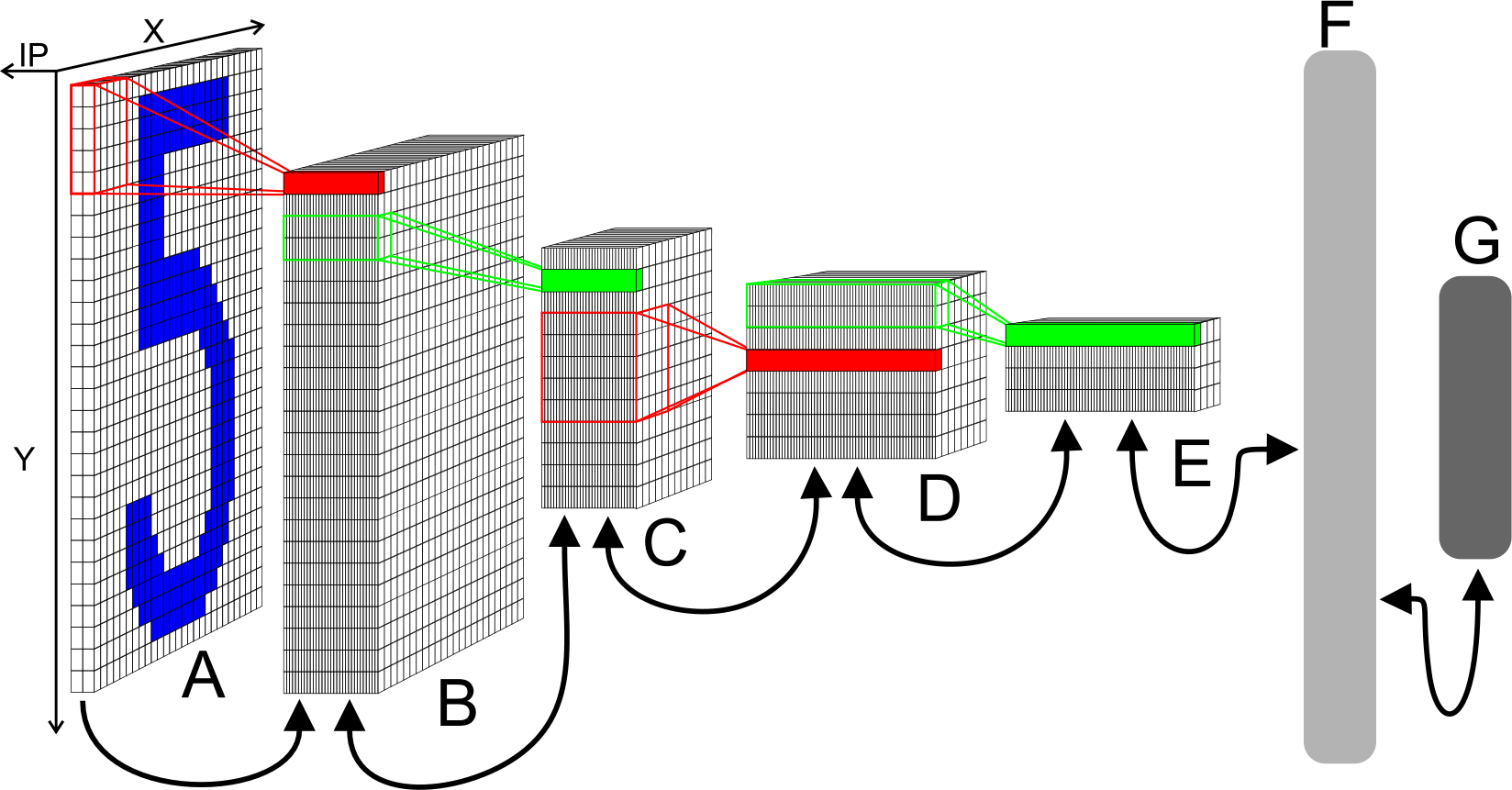
Network structure of the convolution network for MNIST. A: Input layer with 28 × 28 IPs for 28 × 28 input pixel. Each module has two neurons realizing a simplified version of on/off cells. From this layer spikes are send to the layer H1. B: Convolution layer H1 with 24 × 24 IPs with 32 neurons each. Every IP processes the spikes from 5 × 5 blocks of IPs from the input layer (x and y stride is 1) and the spikes from the corresponding pooling cell in layer H2. C: 2 × 2 pooling layer H2 (x and y stride is 2) with 12 × 12 IPs with 32 neurons each. The weights between H1 and H2 are not learned but set to a weight matrix that creates a competition between the 32 features of H1. From convolution layer H3, spikes are processed from IPs with collecting areas that spatial overlap with that H2 IP. Transposed convolution (also known as fractionally strided convolutions) needs to be considered for correctly inverting the shared weights for the weight sets from H2 to H3. D: 5 × 5 convolution layer H3 (x and y stride is 1) with 8 × 8 IPs. Similar to H1 but with 64 neuron for each IP. E: 2 × 2 pooling layer H4 (x and y stride is 2) with 4 × 4 IPs with 64 neurons each. This layer is similar to layer H2. F: Fully connected layer H5. 1024 neurons in one big IP which are fully connected to layer H4 and output layer HY. G: Output layer HY with 10 neurons for the 10 types of digits. For decoding the identity of the output, the neuron with the highest activity is selected.

##### Input layer *X*

The input pattern consists of 28 × 28 pixel with two neurons per pixel after the on/off split. As another computational time saving measure, instead of applying the convolutional kernel to the input spikes and shifting the kernel around, the input is converted into 24 × 24 IPs. Each input population contains the 5 × 5 × 2 neurons that a convolutional kernel would see at that given x and y position on the *H*1 layer. Over these 25 *I*_*On*_(*x, y*) and 25 *I*_*Off*_ (*x, y*) values a normalized probability distribution *p*_*X*_(*s|x, y*) (with *s ∈* [1, 50] and Σ_*s*_*p*_*X*_(*s, x, y*) = 1) is calculated. In every time step of simulation this network, one spike is generated by each of these reorganized input populations. This results in 24 × 24 input spikes per time step in simulation time. These spikes are transmitted to layer *H*1.

##### Convolution layer *H*1

Layer *H*1 receives the spikes produced by input layer *X*. Since the convolution was already realized by re-arranging the input, the weight connections are one to one in the spatial dimensions but uses the same set of weights for all these positions. At every spatial position in *H*1 a IP with 32 neurons is present. Thus the weights *X → H*1 have the dimension (25 × 2) × 32. Not only receives *H*1 spikes from *X*, it also gets spikes from the pooling layer *H*2. During the simulation the latent variables for each IP form a probability distribution *p*_*H*1_(*s|x, y*) = *h*_*H*1_(*s|x, y*) which is used to draw one spike per population in every time step. Thus in every time step 24 × 24 spikes are drawn from *H*1 and send to *H*2. The weights *H*1 *→ H*2 have the dimension (2 × 2 × 32) × 32 due to the network structure that combines a 2 × 2 spatial patch of *H*1 onto one position in *H*2 (with stride 2). Layer *H*1 doesn’t send spikes back to input layer *X*, since the input stays fixed over processing a given input pattern.

##### Pooling layer *H*2

The pooling layer halves the spatial dimensions of layer *H*1 and has 12 × 12 IPs with 32 neurons each. Thus it produces 12 × 12 spikes per time step from the *H*2 latent variables of each IP. Layer *H*2 incorporates the spikes received from a 2 × 2 spatial patch (with stride 2) from layer *H*1 into its latent variables. The weights *H*1 *→ H*2 are designed such that the input from same features at different spatial positions are combined and the 32 features are in a competition. Layer *H*2 sends spikes back to layer *H*1. In this reversed information flow, the spike produced by one spatial position of *H*2 is send to all four connected IPs from the aforementioned 2 × 2 spatial patch (dimensions of weights *H*2 *→ H*1: 32 × (2 × 2 × 32)). This weights can be calculated via 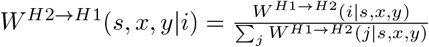. Layer *H*2 sends as well as receives also spikes to / from the next convolution layer *H*3 using a 5 × 5 convolutional kernel (with stride 1). While the forward direction to *H*3 is a normal convolution operation, the backflow of information from *H*3 is a bit more complicated. This operation is called transposed convolution [61] (also known as fractionally strided convolutions). To keep the change of the *H*2 latent variables in every time step, induced by the spikes from *H*3, spatial more uniform, *ϵ*_*Backward*_ is scaled by the number of *H*3 spikes that the corresponding IP in *H*2 receives. A IP in the center of the layer will receive 25 spikes per time step. At the rim, a *H*2 population might only receive 1 spike per time step from *H*3. The dimensions for the weights *H*2 *→ H*3 are (5 × 5 × 32) × 64, since every IP in *H*3 contains 64 neurons. For the other direction *H*3 *→ H*2, the weights have the dimensions 64 × (5 × 5 × 32).

##### Convolution layer *H*3

Layer *H*3 is constructed from 8 × 8 IPs with 64 neurons each which produce 8 × 8 spikes in every time step from their latent variables. The input from *H*2 is processed through a 5 × 5 convolution kernel (with stride 1). The interactions between *H*2 and *H*3 have been described in the paragraph about *H*2 in detail. The information flow between *H*3 *↔ H*4 is like *H*1 *↔ H*2 but with 64 neurons per IP instead.

##### Pooling layer *H*4

Layer *H*4 has 4 × 4 IPs with 64 neurons each. Thus 4 × 4 spikes are produced in every time step and send to layer *H*3 and layer *H*5. Layer *H*5 is a fully connected layer with 1024 neurons. Hence the weights *H*4 *→ H*5 have the dimension (4 × 4 × 64) × 1024 and 1024 × (4 × 4 × 64) for the other direction of information flow. While layer *H*4 sends 16 spikes to layer *H*5 per time step, layer *H*5 sends only one spike back per time step.

##### Fully connected layer *H*5

Layer *H*5 is one big IP with 1024 neurons. It receives 16 spike from layer *H*4 and one spike from output layer *HY* per time step. It send the one spike it generates itself per time step to all neurons of layer *H*4 and *HY*.

##### Output layer *HY*

Layer *HY* is the output layer, which consists of one IP with 10 neurons. In every time step it produces one spike. Every one of these neurons is exclusively associated with one type of digit. For decoding the result of the information processing task, the neuron with the highest *h*_*HY*_ is selected and the digit that is connected to this neuron is used as result.

##### Pre-Learning

Before using the full network, the weights are pre-learned. During pre-learning only two neighboring layers of the network are segregated from the whole network and used for learning as in a two layer network [58] at a time. It needs to be noted that during pre-learning with MNIST, all the input patterns were simplified into black-and-white patterns because this allowed us to speed up the simulation. All the weights are randomly initialized (All the weight values are set to one, 1% uniform positive noise is added and then the weights are normalized). First, the weights between the input layer *X* and the hidden layer *H*1 are batch learned for 20 learning iterations (all these sub-networks are simulated with *ϵ* = 0.1, 1200 spikes per pattern and IP, and *h* is averaged over the last 200 spikes) using the learning rule

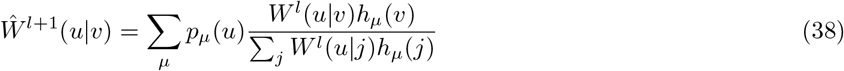

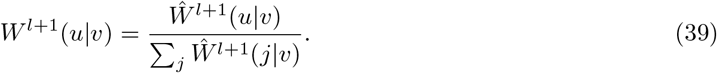

One batch learning step uses the information from all 60,000 training patterns *µ*. *p*_*µ*_(*u*) represents the firing probability distribution of the layer that produces the spikes and *h*_*µ*_(*v*) denotes the hidden layer that processes these spikes. After 20 learning steps, for one more time all patterns are processed for 1200 spikes but this time *h*_*H*1*,µ*_(*v*) is stored as *p*_*H*1*,µ*_(*v*). Then a second two layer network is constructed from hidden layer *H*1 and *H*2. Since the weights for this network are known and don’t need to be learned, *p*_*H*1*,µ*_(*v*) is propagated through this network and results in *p*_*H*2*,µ*_(*v*). Now a third network is created from hidden layer *H*2 and hidden convolutional layer *H*3, using *p*_*H*2*,µ*_(*v*) as input pattern. After 20 learning iterations, *p*_*H*2*,µ*_(*v*) is propagated to *p*_*H*3*,µ*_(*v*) and *p*_*H*3*,µ*_(*v*) is propagated again, using the known pooling weights, into *p*_*H*4*,µ*_(*v*). For the two final sets of weights (*H*4 *→ H*5 and *H*5 *→ HY*) the approach is slightly changed. Like presented in [58], we can convert the remaining three layer network into a two layer network by flipping down the output layer and joining it with layer *H*4 into an extended input layer for this two layer network. During this final pre-learning procedure, 16 spikes from *p*_*H*4*,µ*_(*v*) and one spike from *p*_*HY,µ*_(*v*) are processed by *H*5 in every time step. These two sets of weights are learned for 20 iterations too. After pre-learning, we transposed and re-normalize all the weights produced by pre-learning for also getting the weights for the other direction of information flow.

##### Learning in the full network

The full network (see figure 8) is set up using the pre-learned weights. We use *ϵ*_*Base*_ = 0.4 with 1200 spikes per pattern and IP. We found it helpful to take into account how many spikes are received by a IP and unify the speed of changing for all layers. e.g. one population in *H*3 receives 25 spikes from its connected *H*2 populations, thus we used 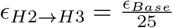. Hence, we used 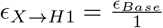, 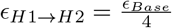, 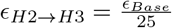, 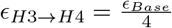, 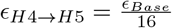, and 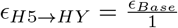 for the direction of information from the input layer *X* to the output layer *HY*. For the other direction of information flow we used 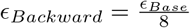 where the receiving population gets one spike as input from this direction. However, we used this 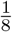 for reducing the influence of the information flowing backwards. The information flow *H*3 *→ H*2 is an exception because *ϵ*_*Backward*_ is scaled down by the amount of spikes the receiving populations in *H*2 get, too. How strongly *ϵ* is reduced depends, due to the transposed convolution, on the spatial position of the *H*2 IP. For learning, the same learning rule from pre-learning is used. Furthermore, only the weights for *X → H*1, *H*2 *→ H*3, *H*4 *→ H*5, and *H*5 *→ HY* are learned actively. The weights for the other direction are calculated from them again. Before each new learning iteration commences, a small offset (*max*(*W* (*u|V*))*/N* with *N* as total number of matrix elements) is added to the weights for preventing the multiplicative learning rule to get stuck.

In figure 9, the classification performance is shown for six learning steps of the full network. At learning step zero, we used the weights from pre-learning. We tested the performance for the full network with feedback, as used for learning, and also with setting *ϵ*_*Backward*_ = 0. Converting the aforementioned Tensor Flow tutorial network to this structure ends in a 2.9% error. We reach with the SbS network a minimum of 2.2% error. However, the performance of the Tensor Flow network drastically increased (over 99% correct classification) if more advanced learning strategies (e.g. Adam [60] or L4 [62] and input distortion methods) are used and not only a simple gradient method. It needs to be noted that due to a lack of computational power, we couldn’t even try to optimize the parameters of the SbS network. We guessed a set of parameters and used it. For this reason we also only used our standard batch learning rule. No tests, while tempting, with ADAM or L4 derivatives were possible. The remaining pictures concerning the MNIST dataset use the weights from the 5th learning step. Figure 9 shows that using local learning rules with information flowing from the output layer to the input layer only via spikes is, even in a deep network, possible.

**Fig 9.**
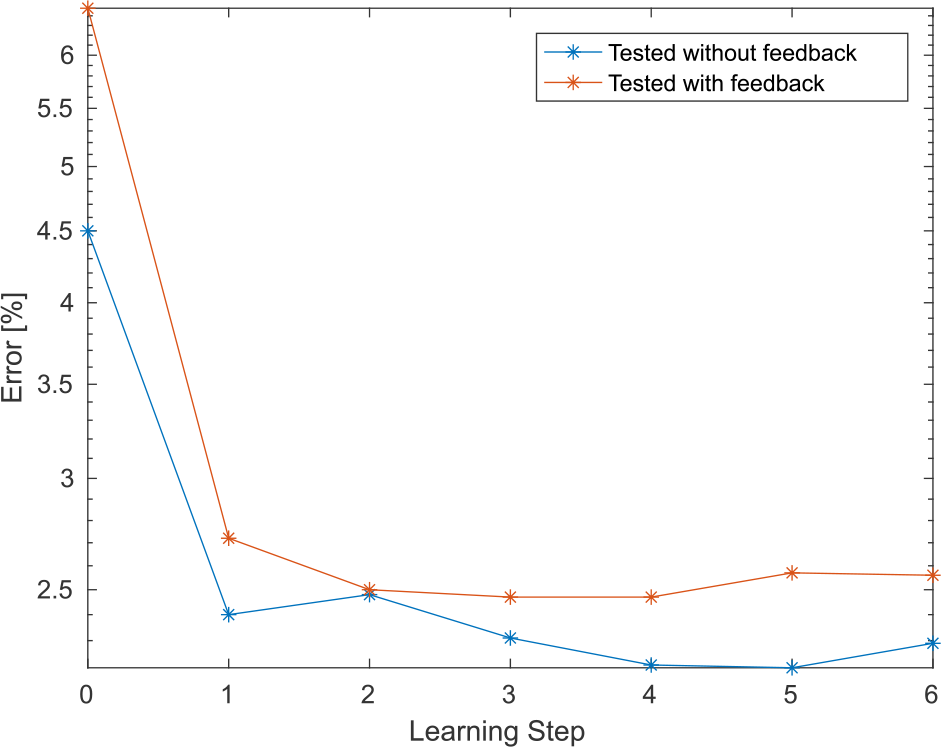
Performance of the convolution network on the MNIST benchmark. First the weights of the network are pre-learned with the batch learning rule in a mostly unsupervised fashion. At this point the network shows an error on the test dataset of 6.48% with active feedback and 4.5% without feedback spikes. Then the weights are subjected to 6 learning steps with batch learning in the complete network with active feedback for fine-adjustment of the pre-learned weights. The red line shows the test data error with active feedback and the blue line shows the test data without feedback. After 5 learning steps, the error reaches a minimum of 2.2% for the test data without feedback.

##### Receptive fields

In figure 10 (layer *H*1), figure 11 (layer *H*3), figure 12 (layer *H*5), and figure 13 (output layer *HY*) the receptive fields (RFs) of parts of the network are shown. We used two types of inputs for visualizing the RFs: a.) 500,000 random (uniform noise) image pattern with 28 × 28 were generated and processed (for 1200 spike per pattern and IP) by the network. b.) The 60,000 training patterns and 10,000 test patterns were processed by the network. After the network has processed the spikes, then the resulting *h*_*µ*_ are stored. The RFs are simply calculated via 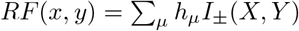.

**Fig 10.**
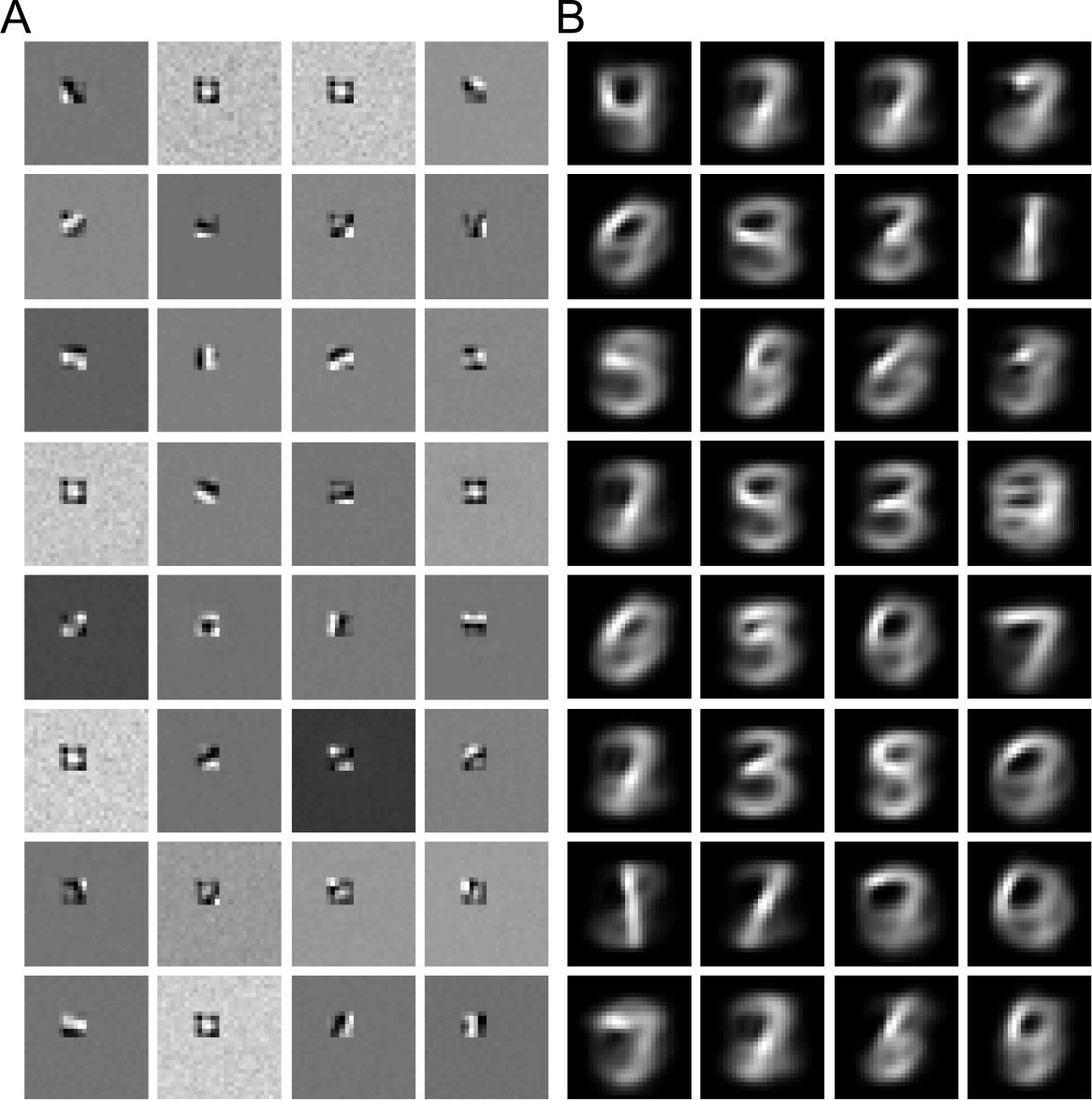
MNIST Network: Reversed correlation for convolution layer. *H*1. Reverse correlation in the MNIST network (with feedback) of the IP at the spatial coordinate (10,10) with a: random noise and b: with the test and training data set. Results for pooling layer *H*2 look very similar, thus not show.

**Fig 11.**
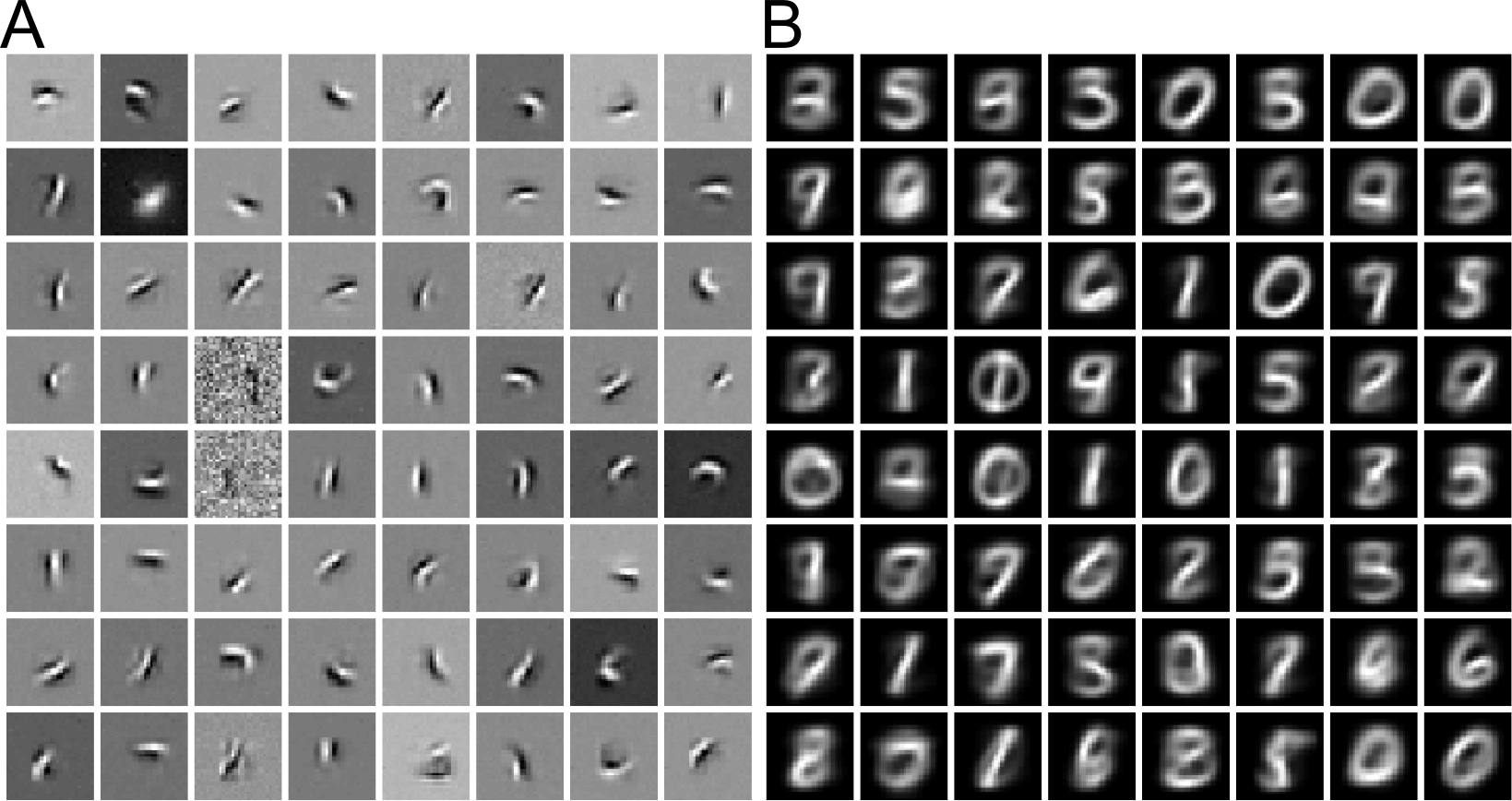
MNIST Network: Reversed correlation for convolution layer. *H*3. Reverse correlation of the IP at the spatial coordinate (5,5) with a: random noise and b: with the test and training data set. Results for pooling layer *H*4 look very similar, thus not show.

**Fig 12.**
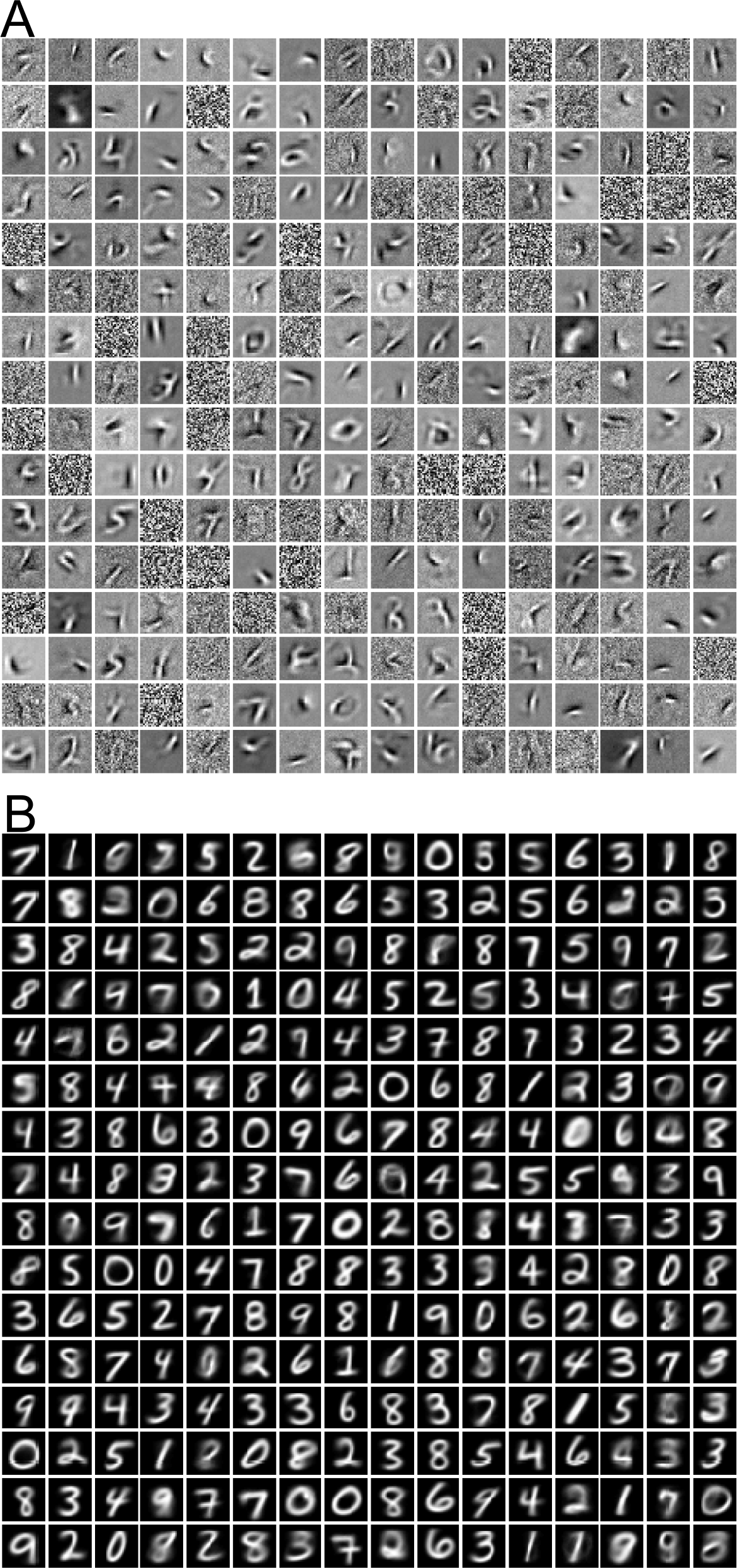
MNIST Network: Reversed correlation for fully connected layer. *H*5. Reverse correlation shown for 256 neurons of the 1024 neurons with a: random noise and b: with the test and training data set.

**Fig 13.**
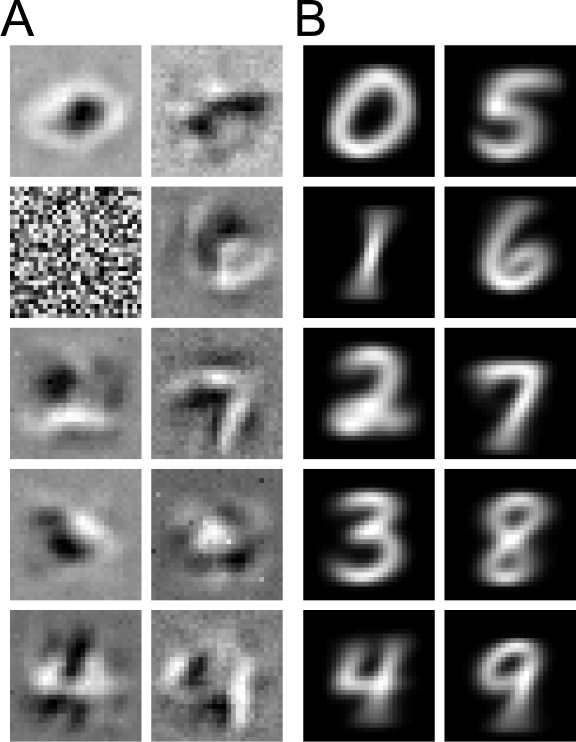
MNIST Network: Reversed correlation for fully connected output layer*HY*. Reverse correlation shown for the output neurons with a: random noise and b: with the test and training data set.

While the RFs calculated from the random patterns show what is locally interesting for the neuron, the RFs based on the MNIST data shows how the data statistics extends it to the whole input field. The responses of the network to the random input pattern is rather weak. The pooling layers are not shown, because the RFs look rather similar to the ones from the layer before but spatially extended due to the pooling operation. In the figures, every tile shown is scaled independently to a [0, 1] value range for improving visibility. The closer the layer gets to the output layer, the more complex the RFs gets or more distinctly look like a digit in the case with the non-random input patterns.

##### Generative model

While the SbS model is inspired by generative models [58], it is interesting to see what happens to is generative property in a deep network. Figure 14 shows an example how the *h*-distribution of layer *H*1 can be used to reconstruct the input pattern from it. While this reconstruction resembles the input very well, using the activity of the higher layers leads to a disastrous reconstitution. The reason for this lies in that the network was optimized for processing information (e.g. estimating the type of digit from the input) and not representing the input as best as possible. The pooling layers have also a very destructive effect in this regard because they spatially smear information.

**Fig 14.**
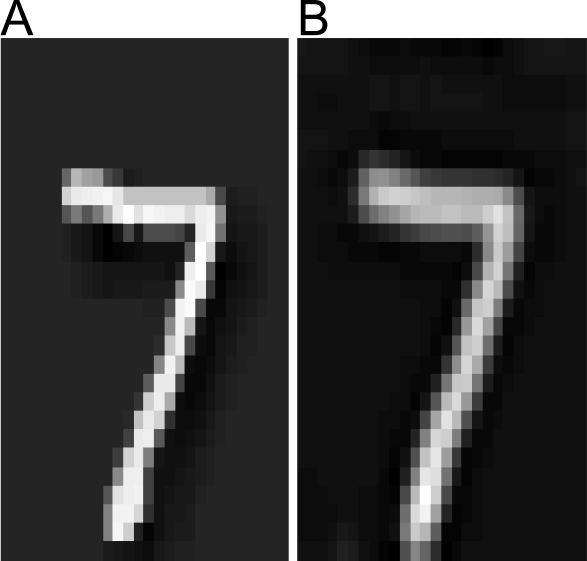
MNIST Network: Example of reconstructing the input from the activity of layer. *H*1. a: This example image of a 7 was feed into the MNIST network and 1200 spikes per IP were processed. b: Based on the resulting *H*1 activities, a reconstruction of the input was calculated.

### Natural images

In SbS networks learning corresponds to finding weights that will maximize the harmony of latent variable configurations in the network averaged over the ensemble of inputs. This is formally independent from optimizing particular input-output functions and therefore represents a form of self-organization. The framework therefore might have some potential of serving as a model for cortical function where supervised learning of the connectivities is implausible. We therefore investigated if unsupervised learning with natural images of a network with the architecture used for the above MNIST data classification could reproduce properties of the visual cortex that depend on the feedback from deeper layers.

Removing the output layer *HY* from the structure of the SbS MNIST network (shown in figure 8), we trained the network with 28 × 28 gray natural image patches. We generated these image patches from the full image McGill calibrated color image database http://tabby.vision.mcgill.ca/. We took the high resolution color images, used the rgb2linear function, and collapsed the color channels into gray values. We whitened the picture via a singular value decomposition method and cut out 28 × 28 patches at random positions (avoiding a dead pixel from the camera). For learning, first the pre-learning procedure was applied for 30 learning steps each, using the gray values as input. Then the full network learned the weights for another 5 learning steps. For every learning step performed (pre-learning and full network), 50,000 new input images were generated and used as training data.

Figure 15, figure 16, and figure 17 show RFs for the layers *H*1, *H*3 and *H*5. As input pattern 500,000 random images were used as well as the 1.75 million training patterns. In these figures, every tile shown is scaled independently to a [0, 1] value range for improving visibility. The further the neuron is away from the input, the more structured and spatially extended the RF gets, as is the case in the visual system.

**Fig 15.**
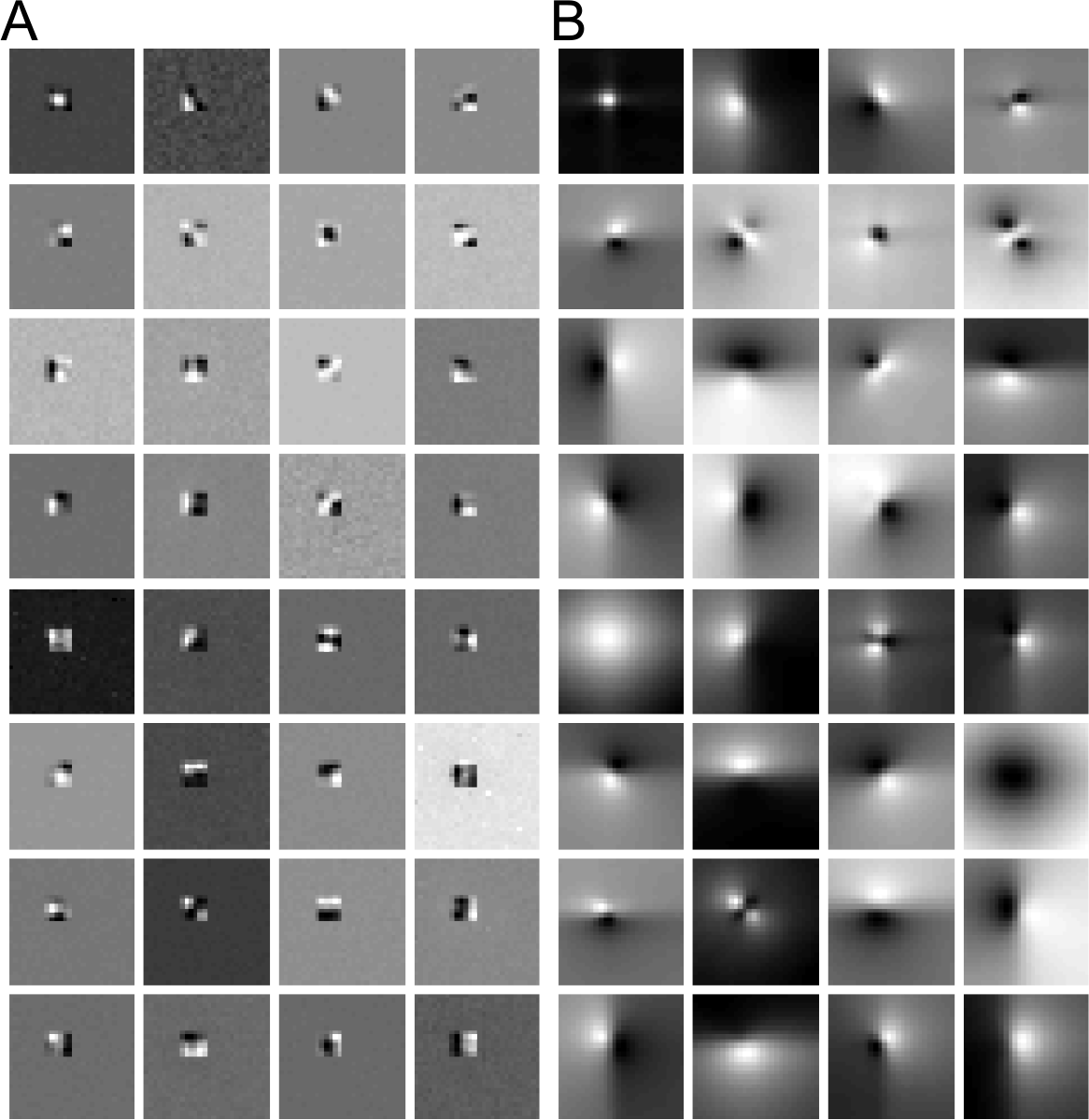
Natural image network: Reversed correlation for convolution layer. *H*1. Reverse correlation in the natural image network (with feedback; like the MNIST network shown in Figure 8 but without the output layer *HY*) of the IP at the spatial coordinate (10,10) with a: random noise and b: with the training data set. Results for pooling layer *H*2 look very similar, thus not show.

**Fig 16.**
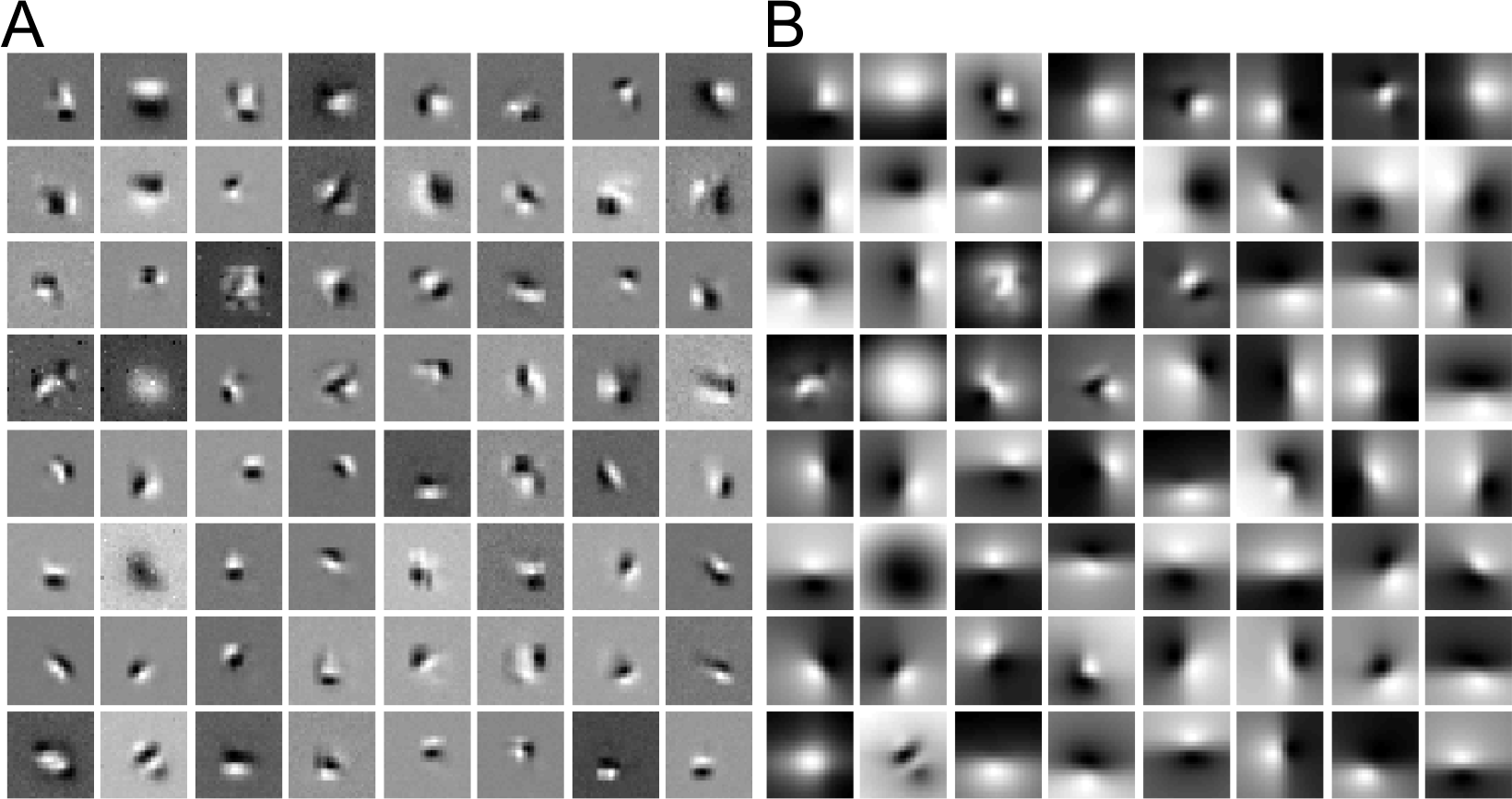
Natural image network: Reversed correlation for convolution layer. *H*3. Reverse correlation of the IP at the spatial coordinate (5,5) with a: random noise and b: with the training data set. Results for pooling layer *H*4 look very similar, thus not show.

**Fig 17.**
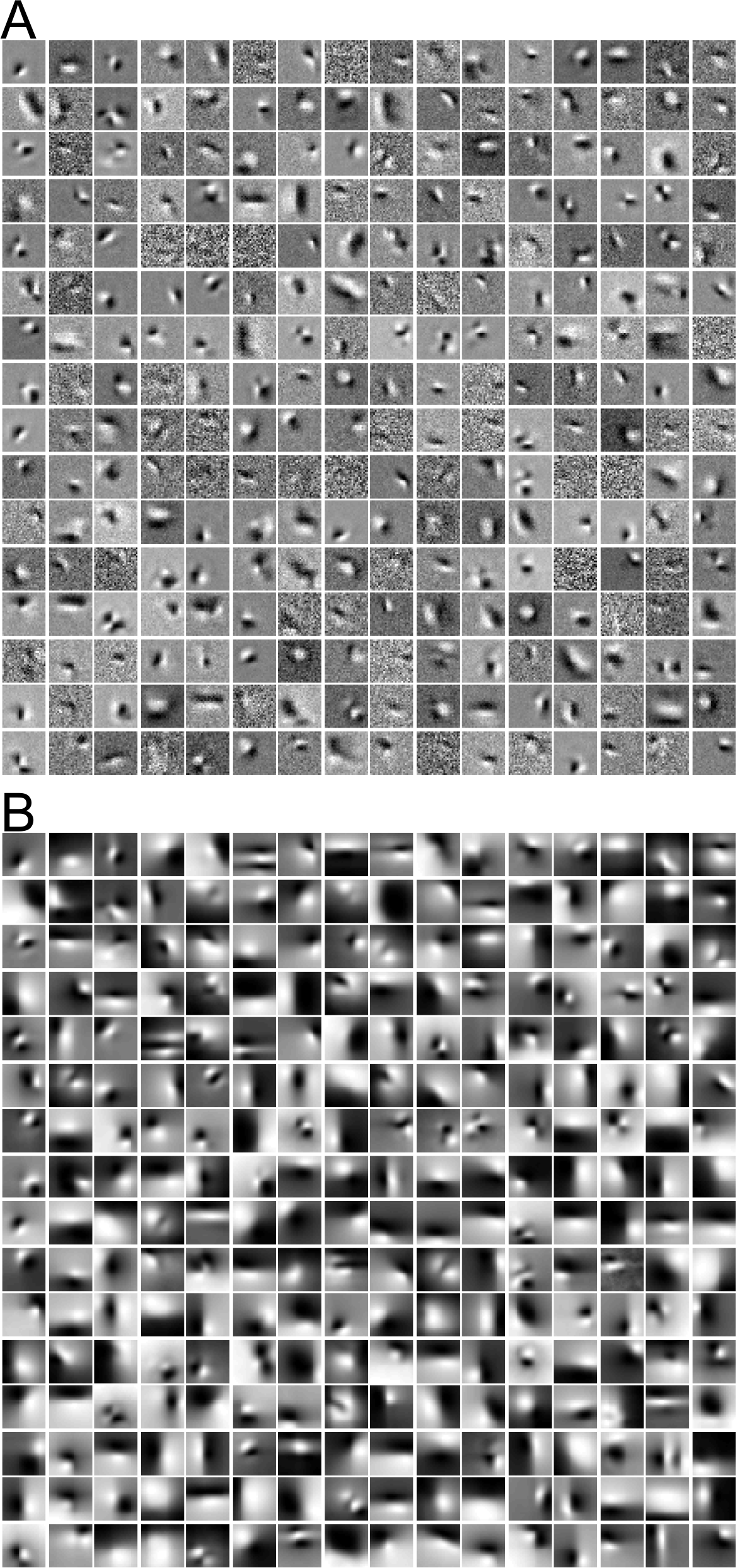
Natural image network: Reversed correlation for fully connected layer *H*5. Reverse correlation shown for 256 neurons of the 1024 neurons with a: random noise and b: with the training data set.

The forward architecture of this model is structurally similar to DCNs that were used for the purpose of explaining response properties of neurons in visual cortex [63]. Since here the deeper layers provide feedback input to their respective previous layers, we checked if this model might reproduce effects in cortex that are known to depend on recurrent interactions from deeper layers [64].

We tested the relevance of these RFs for linear prediction of the neuron’s responses. To this end, we took the RFs and 50,000 input pattern, removed their means and normalized both, the RFs and the input patterns according the L2 norm over their spatial dimension. This results in unity vectors for the receptive field 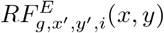 (with *g* representing the layer, *x′* and *y′* the spatial postion of the IP and *i* the neuron in the IP) as well as 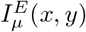 for the input pattern. The scalar product between these vectors is calculated:

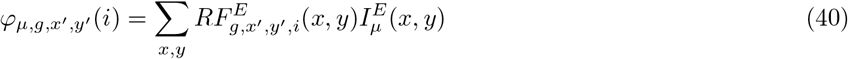

On the other side we have the latent variables *h*_*µ,g,x′,y′*_(*i*) as the asymptotic result of processing the input patterns *I*_*µ*_(*x, y*) by the whole network. After removing their mean and dividing by their standard deviation, we get 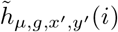 and 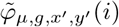. Using

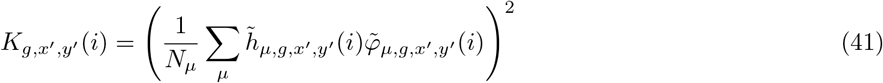

we calculate the correlation of these two entities for each input pattern. Averaging *K*_*g,x′,y′*_(*i*) over *x′*, *y′* and *i* yields 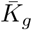.

Using the RFs obtained with noise patterns, this results in 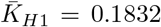, 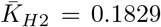,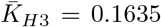, 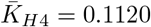, and 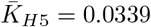, respectively, for the network with feedback and 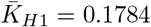, 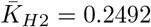, 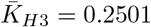, 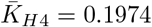, and 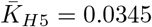 for networks where the feedback was disabled. Using the RFs generated from the natural images, we obtain 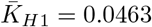, 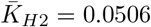, 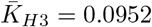, 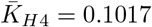, and 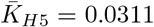, respectively, for the network with feedback and 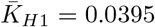, 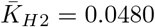, 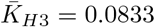, 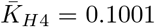, and 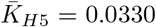 without feedback. The lower values for the case using the RFs calculated from the natural images is a result of the structured surround area in RFs as a result from the data statistics in the natural image input patterns.

In summary, for layers close to the input the RFs can explain a substantial part of the neuron’s activities with a linear model, which in visual cortex would correspond to simple cells in V1. The responses in the deeper layers turn out to be more nonlinear which would in cortex correspond to more complex responses downstream in the visual system.

Then we investigated the quality of coding of the input. Based on the *H*1 activities after presenting 50,000 natural images, we reconstructed 5×5 pixel input pattern patches *I*_*µ,x′,y′*_(*x, y*) for the IP at the spatial positions *x′* and *y′*. Corresponding to its *H*1 neurons we reconstructed 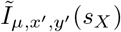 according to

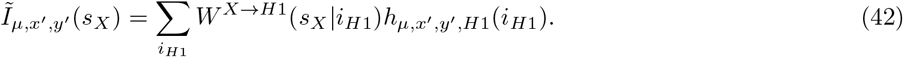

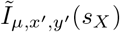 is then transformed, by recombining the on/off channels in one gray value channel, into 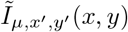 (which lives normal pixel space). From 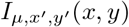 and 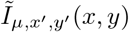 the respective average mean over all 25 pixels is calculated and subtracted. Then 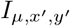 and 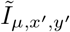are normalized via the L2-norm. Thus we get unit vectors 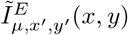 and 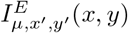. The scalar product between these two types of vectors are calculated:

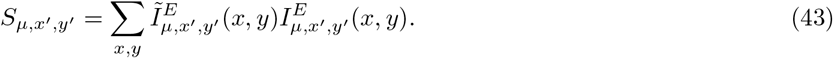

Calculating over all possible 24×24 *x′* and *y′* positions as well as the 50,000 pattern, results in 0.550 *±* 8 *⋅* 10^*−*5^ (std error of the mean) with feedback and 0.537 *±* 8 *⋅* 10^*−*5^ (std error of the mean) without feedback. The less than perfect quality of the reconstruction is a result of using the same convolution weights for all positions of input space. Thus the weights are a compromise for all spatial positions.

We wondered if already this toy model might reproduce effects in primary visual cortex that depend on the feedback from higher areas. For this purpose we compared the responses of neurons in the first layer *H*1 when stimulated with their own RF obtained from natural images (Figure 15 b) with the case where the feedback connections where disabled (Fig. 18). Strikingly, we find that the majority of neurons exhibit higher activity when feedback was switched off. This result matches the recent experimental finding [64], that the feedback from area V2 contributes substantially to the surround suppression of responses in area V1.

**Fig 18.**
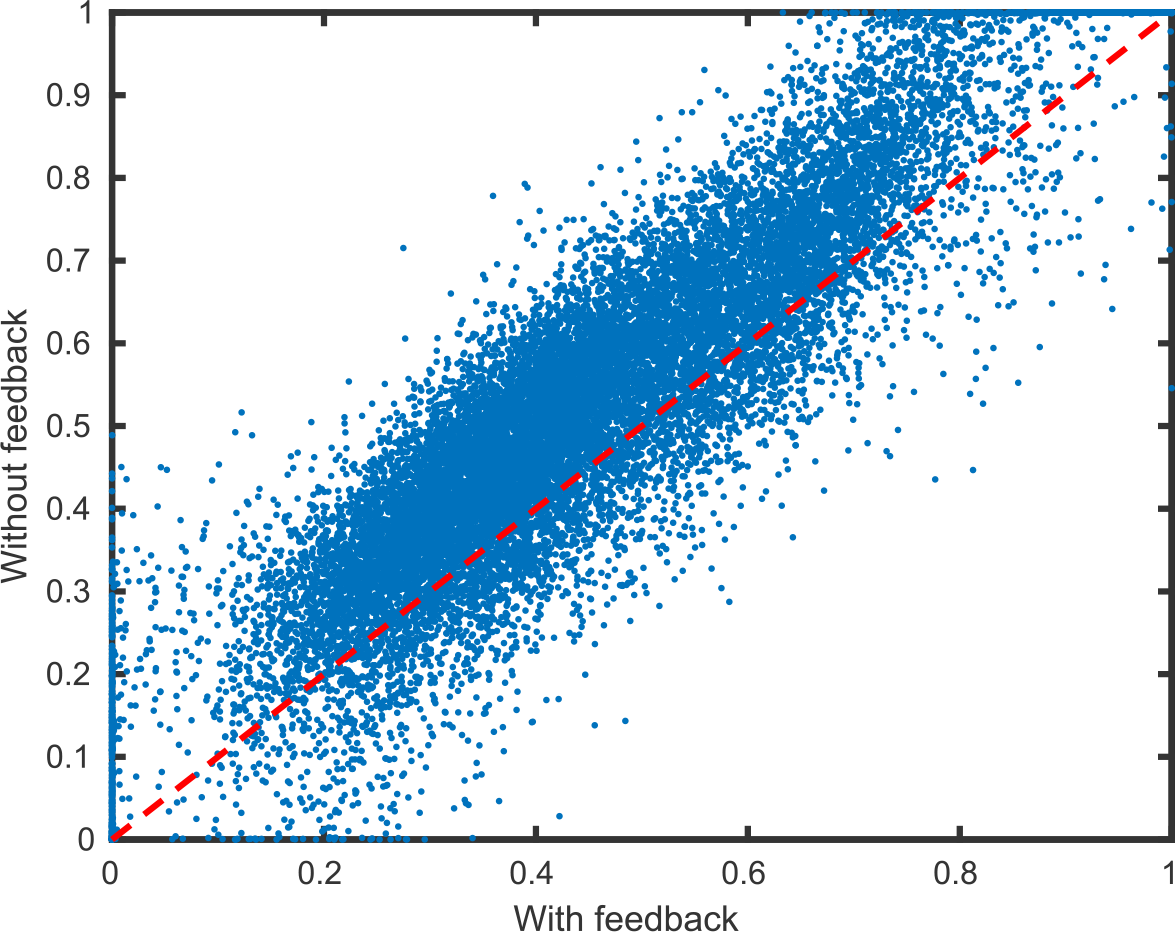
Influence of the feedback connection on the activity of *H*1. Using the shown RF in figure 15b of the *H*1 neurons (calculated from the natural images) as input for the network with and without active feedback. The figure shows all the 18432 *H*1 neurons reacting to their own RF with and without feedback from the higher layers.

## Discussion

Deep Convolutional Networks (DNCs) are particularly successful artificial neural networks [4]. However, they lack biological realism which sows doubt that they can be considered also as models for real neuronal computations. The current work presents a framework with competitive computational abilities for large neuronal networks with virtually arbitrary architectures that employs biologically plausible mechanisms.

The nodes in this novel approach represent sparse coding neuronal circuits (SCNC) instead of single neurons. Each SCNC encompasses a population of neurons which jointly develop a sparse representation of all spikes it receives as input. Originally, such circuits were introduced for explaining the response properties of simple cells in visual cortex [44]. Meanwhile the principle of sparse neuronal coding has been related to compressed sensing [65], a general coding principle that might explain the function of a wide range of neuronal systems [43, 49].

There are several detailed models for such circuits that employ spiking neurons [47, 48, 56]. For the present work, however, we skip detailed modeling of the circuits and replace them by a minimal model – the inference population (IP) – that mimics the neuronal dynamics leading to sparse efficient coding with each impinging spike [58]. The latent variable of every neuron in a IP is updated with each spike received by that IP to which the neuron belongs. It reflects internal states of the neuron which determine its activity as e.g. the membrane potential. We identify it with the firing rate of the neuron and use it to generate spikes as means of communicating with other populations. In particular, the framework is consistent with spikes being generated according to independent Poisson point processes. This mimics the statistics of neuronal activities in cortex [66]. In fact, the stochasticity of spiking in cortex is believed to reflect the balance of excitation and inhibition within local circuits [67, 68] and has been reproduced in more detailed models of SCNCs [69].

In contrast to realistic models of neuronal circuits, our IP model for the SCNC skips the computations during the time in between spikes. Besides formal tractability, this spike-by-spike update dynamics is computationally far more efficient than more bio-physically realistic models [70]. Thus, networks of IPs (NIPs) present an interesting alternative situated between the biologically unrealistic DCNs and networks with more biologically realistic neuron models as e.g. leaky integrate-and-fire neurons.

The dynamics of the latent variables corresponds to an optimization of the representation of the respective inputs of each circuit. We found that this induces also an optimization of a global objective function for the whole network that reflects a global coherence of all activities. While this might be seen as a specific realization of the ‘Free Energy Principle’ [71], we instead termed this objective ‘harmony’.

The idea to learn hierarchical networks composed of auto-encoders in a biologically realistic way has been sketched before [72]. To our knowledge, however, the field of generative networks has not yet proposed a framework that combines the features of the present approach: it allows for arbitrary architectures, uses realistic interactions among the circuits, and obeys essential neurobiological constraints. In particular, each circuit in our framework sends and processes only stochastic spikes. Also, learning in the present framework corresponds to changing only non-negative inter-circuit synaptic efficacies. Thereby it keeps the weights on these long-range excitatory connections excitatory, which is the case in cortex (Dale’s law).

[58] already showed that one large IPs can learn and realize arbitrary Boolean functions by representing each entry in the truth panel. Here, we use the example of parity functions to demonstrate that a NIPs can solve the same task far more efficiently in a multi-layer hierarchical fashion. This shows that the framework generates sufficiently strong non-linearities for performing arbitrary computations and hints at its generality along the lines of the pioneering work of McCullogh and Pitts [1].

The learning rule is derived from the same objective function termed ‘harmony’ as used for obtaining the IP’s dynamics. Reformulating this learning rule in terms of the dynamics of the neuronal activities reveals that it consists of two terms, one representing a differential Hebb rule, and one corresponding to the ordinary Hebbian rule. Also, the weights are post-synaptically regulated by normalization. This combination corresponds to a specific form of Spike-Timing Dependent Plasticity [25–27].

For supervised learning the targets are simply presented as additional inputs to the output layer. For the special case of hierarchical architectures analog to those in Deep Convolutional Networks used for pattern recognition we found that this approach is sufficient for learning Boolean functions as well as for training deep networks for pattern recognition. Application to the MNIST benchmark data http://yann.lecun.com/exdb/mnist are used to demonstrate that the approach can achieve substantial performances that are comparable to those of DCNs when using identical network architectures, applying simple gradient-optimization methods, and using the unaltered training data set for both networks. In the case of the DCN a error back-propagation learning rule is applied, while the IPs use only a local learning rule. Interestingly, networks of IPs that have only feed-foreward architectures can also be optimized by a rule similar to back-propagation [57], where we obtain very similar performances. This further underlines the efficiency of learning by optimizing the ‘harmony’.

While for learning by optimizing the ‘harmony’ the feedback connections are essential, they can be a cause for performance reduction during classification. The good performances for the toy example of digit classification are encouraging, however, the IP based networks can not yet compete with DCNs, when for the DCNs state-of-the-art learning methods are applied (e.g. [60, 62]) and the size of the training data set is increased by methods of input pattern distortion. The latter we couldn’t apply due to restriction on available computation power and the large effort that would be required for searching for optimal parameters (e.g. searching for optimized *ϵ* values). The former reveals a lack of a good learning procedure for technical applications with NIPs – which in comparison had be developed and improved over many years for DCNs –, which will be an interesting research question for the future. For the latter, the development of specialized hardware contributed substantially to the success of DCNs [73, 74] since it allowed for highly efficient implementations of large networks. Also for NIPs extensive simulations will also become possible from the fact that the IPs in this framework are local circuits that can operate independently and in parallel. This will allow to build special hardware [75] that in future can be used in technical applications as well as large scale models of networks in the brain.

The feedback connections can serve to complement missing inputs and for realizing context influences. Already the example of the Boolean parity functions demonstrates that context effects can be dramatic, where a single bit decides about the computation for all other inputs. Generally, depending on particular inputs, a given network’s function can be switched depending on signals that represent e.g. the state of attention or task. This demonstrates that the ‘harmony’ might be a principle that serves to control a network’s function and shows that one does not need to assume special mechanisms for realizing context effects or attentional reconfiguration. On the other hand the objective of finding a configuration of latent variables that maximizes the consistency bears the danger of ignoring aspects of the input. While this effect might underlies visual illusions and hallucinations it on the other side makes computations robust against missing, distorted and occluded inputs, which is a hallmark of natural cognitive systems.

The framework allows for arbitrary network architectures as e.g. in the visual system where the areas are recurrently connected [76] and where no simple hierarchy is present which would allow for a clear cut distinction between ‘bottom up’ and ‘top down’ processing. Also, it can provide a spike based modeling approach for multi-sensory integration [77].

As a first step towards modeling cortical networks, we adopted the architecture of the network for classification of the hand-written digits, however, we removed the last layer and then trained the remaining network with natural images. In the first layer this self-organization yields receptive fields similar to the primary visual cortex and expected from previous work with single SCNCs [40, 45]. Responses in the deeper layers turned out to be increasingly more nonlinear which more and more prohibits a characterization as receptive fields of simple cells. Most importantly, the feedback connections predominantly have a suppressive effect for stimuli extending the classical receptive field. These contributions to non-classical receptive field properties match recent results in area V1 of visual cortex [64] that by construction cannot be captured by DCNs. We speculate that with architectures more similar to the conditions of the visual system, our framework will explain also other context dependencies of responses.

The vision that coding and computations in the brain might be understood within the framework of generative models dates back to Heinrich von Helmholtz [78]. Since then it remained influential in many aspects of brain science. More recently, generative models also came into the focus of research in machine learning and had impressive successes [79]. While the proposed NIPs fall into the class of generative models, however, we feel it is far too early for speculations that the current approach could explain relevant aspects of coding, computation, and learning in the brain. Most importantly, in its current form it is restricted to mainly static inputs which are not the most relevant stimuli in reality. As a first step towards the goal of enabling also temporal processing within this realistic approach, however, it could be suffcient to include delay lines, which we will explore in our future research.

## Acknowledgments

We thank Udo Ernst fruitfull discussions. This work was supported in part by Bundesministerium fuer Bildung und Forschung Grant 01 GQ 1106 (Bernstein Award Udo Ernst) as well as the Creative Unit I-See ‘The artifical eye: Chronic wireless interface to the visual cortex’ at the University of Bremen. A patent was filed.

## References

1. McCulloch WS, Pitts W. A logical calculus of the ideas immanent in nervous activity. The bulletin of mathematical biophysics. 1943;5(4):115–133.

2. Ackley DH, Hinton GE, Sejnowski TJ. A learning algorithm for Boltzmann machines. Cognitive science. 1985;9(1):147–169.

3. Dayan P, Hinton GE, Neal RM, Zemel RS. The helmholtz machine. Neural computation. 1995;7(5):889–904.

4. LeCun Y, Bengio Y, Hinton G. Deep learning. nature. 2015;521(7553):436.

5. Schmidhuber J. Deep learning in neural networks: An overview. Neural networks. 2015;61:85–117.

6. Guo Y, Liu Y, Oerlemans A, Lao S, Wu S, Lew MS. Deep learning for visual understanding: A review. Neurocomputing. 2016;187:27–48.

7. Azkarate Saiz A. Deep learning review and its applications; 2015.

8. Silver D, Huang A, Maddison CJ, Guez A, Sifre L, Van Den Driessche G, et al. Mastering the game of Go with deep neural networks and tree search. nature. 2016;529(7587):484.

9. Mnih V, Kavukcuoglu K, Silver D, Rusu AA, Veness J, Bellemare MG, et al. Human-level control through deep reinforcement learning. Nature. 2015;518(7540):529.

10. Gatys LA, Ecker AS, Bethge M. Image style transfer using convolutional neural networks. In: Proceedings of the IEEE Conference on Computer Vision and Pattern Recognition; 2016. p. 2414–2423.

11. Längkvist M, Karlsson L, Loutfi A. A review of unsupervised feature learning and deep learning for time-series modeling. Pattern Recognition Letters. 2014;42:11–24.

12. Shen D, Wu G, Suk HI. Deep learning in medical image analysis. Annual review of biomedical engineering. 2017;19:221–248.

13. Angermueller C, Pärnamaa T, Parts L, Stegle O. Deep learning for computational biology. Molecular systems biology. 2016;12(7):878.

14. Kriegeskorte N. Deep neural networks: a new framework for modeling biological vision and brain information processing. Annual review of vision science. 2015;1:417–446.

15. Li Y. Deep reinforcement learning: An overview. arXiv preprint arXiv:170107274. 2017;.

16. Miotto R, Wang F, Wang S, Jiang X, Dudley JT. Deep learning for healthcare: review, opportunities and challenges. Briefings in bioinformatics. 2017;19(6):1236–1246.

17. Rawat W, Wang Z. Deep convolutional neural networks for image classification: A comprehensive review. Neural computation. 2017;29(9):2352–2449.

18. Aliper A, Plis S, Artemov A, Ulloa A, Mamoshina P, Zhavoronkov A. Deep learning applications for predicting pharmacological properties of drugs and drug repurposing using transcriptomic data. Molecular pharmaceutics. 2016;13(7):2524–2530.

19. McCann MT, Jin KH, Unser M. Convolutional neural networks for inverse problems in imaging: A review. IEEE Signal Processing Magazine. 2017;34(6):85–95.

20. Marblestone AH, Wayne G, Kording KP. Toward an integration of deep learning and neuroscience. Frontiers in computational neuroscience. 2016;10:94.

21. Kheradpisheh SR, Ghodrati M, Ganjtabesh M, Masquelier T. Deep networks can resemble human feed-forward vision in invariant object recognition. Scientific reports. 2016;6:32672.

22. VanRullen R. Perception science in the age of deep neural networks. Frontiers in psychology. 2017;8:142.

23. Rajalingham R, Issa EB, Bashivan P, Kar K, Schmidt K, DiCarlo JJ. Large-scale, high-resolution comparison of the core visual object recognition behavior of humans, monkeys, and state-of-the-art deep artificial neural networks. Journal of Neuroscience. 2018;38(33):7255–7269.

24. Kietzmann TC, McClure P, Kriegeskorte N. Deep neural networks in computational neuroscience. bioRxiv. 2018; p. 133504.

25. Levy W, Steward O. Temporal contiguity requirements for long-term associative potentiation/depression in the hippocampus. Neuroscience. 1983;8(4):791–797.

26. Markram H, Lübke J, Frotscher M, Sakmann B. Regulation of synaptic efficacy by coincidence of postsynaptic APs and EPSPs. Science. 1997;275(5297):213–215.

27. Bi Gq, Poo Mm. Synaptic modifications in cultured hippocampal neurons: dependence on spike timing, synaptic strength, and postsynaptic cell type. Journal of neuroscience. 1998;18(24):10464–10472.

28. Rumelhart DE, Hinton GE, Williams RJ. Learning representations by back-propagating errors. nature. 1986;323(6088):533.

29. Bengio Y, Laufer E, Alain G, Yosinski J. Deep generative stochastic networks trainable by backprop. In: International Conference on Machine Learning; 2014. p. 226–234.

30. Lee JH, Delbruck T, Pfeiffer M. Training Deep Spiking Neural Networks Using Backpropagation. Frontiers in Neuroscience. 2016;10:508. doi:10.3389/fnins.2016.00508.

31. Anwani N, Rajendran B. Training Multilayer Spiking Neural Networks using NormAD based Spatio-Temporal Error Backpropagation. arXiv preprint arXiv:181110678. 2018;.

32. Wu Y, Deng L, Li G, Zhu J, Shi L. Spatio-temporal backpropagation for training high-performance spiking neural networks. Frontiers in neuroscience. 2018;12.

33. Hinton GE. A practical guide to training restricted Boltzmann machines. In: Neural networks: Tricks of the trade. Springer; 2012. p. 599–619.

34. Salakhutdinov R, Hinton G. Deep boltzmann machines. In: Artificial intelligence and statistics; 2009. p. 448–455.

35. Sountsov P, Miller P. Spiking neuron network Helmholtz machine. Frontiers in computational neuroscience. 2015;9:46.

36. Liu W, Wang Z, Liu X, Zeng N, Liu Y, Alsaadi FE. A survey of deep neural network architectures and their applications. Neurocomputing. 2017;234:11–26.

37. Hochreiter S, Schmidhuber J. Long short-term memory. Neural computation. 1997;9(8):1735–1780.

38. Dayan P. Helmholtz machines and wake-sleep learning. Handbook of Brain Theory and Neural Network MIT Press, Cambridge, MA. 2000;44(0).

39. Douglas RJ, Martin KA. Opening the grey box. Trends in Neurosciences. 1991;14(7):286–293.

40. Lee DD, Seung HS. Learning the parts of objects by non-negative matrix factorization. Nature. 1999;401(6755):788.

41. Lee DD, Seung HS. Algorithms for non-negative matrix factorization. In: Advances in neural information processing systems; 2001. p. 556–562.

42. Salakhutdinov R. Learning deep generative models. Annual Review of Statistics and Its Application. 2015;2:361–385.

43. Ganguli S, Sompolinsky H. Compressed sensing, sparsity, and dimensionality in neuronal information processing and data analysis. Annual review of neuroscience. 2012;35:485–508.

44. Olshausen BA, Field DJ. Emergence of simple-cell receptive field properties by learning a sparse code for natural images. Nature. 1996;381(6583):607.

45. Olshausen BA, Field DJ. What is the other 85 percent of V1 doing. L van Hemmen, & T Sejnowski (Eds). 2006;23:182–211.

46. Rozell CJ, Johnson DH, Baraniuk RG, Olshausen BA. Sparse coding via thresholding and local competition in neural circuits. Neural computation. 2008;20(10):2526–2563.

47. Moreno-Bote R, Drugowitsch J. Causal inference and explaining away in a spiking network. Scientific reports. 2015;5:17531.

48. Zhu M, Rozell CJ. Modeling inhibitory interneurons in efficient sensory coding models. PLoS computational biology. 2015;11(7):e1004353.

49. Ganguli S, Sompolinsky H. Statistical mechanics of compressed sensing. Physical review letters. 2010;104(18):188701.

50. Spanne A, Jörntell H. Questioning the role of sparse coding in the brain. Trends in neurosciences. 2015;38(7):417–427.

51. Palm G. Neural associative memories and sparse coding. Neural Networks. 2013;37:165–171.

52. King PD, Zylberberg J, DeWeese MR. Inhibitory interneurons decorrelate excitatory cells to drive sparse code formation in a spiking model of V1. Journal of Neuroscience. 2013;33(13):5475–5485.

53. Babadi B, Sompolinsky H. Sparseness and expansion in sensory representations. Neuron. 2014;83(5):1213–1226.

54. Balavoine A, Romberg J, Rozell CJ. Convergence and rate analysis of neural networks for sparse approximation. IEEE transactions on neural networks and learning systems. 2012;23(9):1377–1389.

55. Shapero S, Zhu M, Hasler J, Rozell C. Optimal sparse approximation with integrate and fire neurons. International journal of neural systems. 2014;24(05):1440001.

56. Zhu M, Rozell CJ. Visual nonclassical receptive field effects emerge from sparse coding in a dynamical system. PLoS computational biology. 2013;9(8):e1003191.

57. Rotermund D, Pawelzik KR. Back-propagation learning in deep Spike-By-Spike networks. bioRxiv. 2019;doi:10.1101/569236.

58. Ernst U, Rotermund D, Pawelzik K. Efficient computation based on stochastic spikes. Neural computation. 2007;19(5):1313–1343.

59. Roberts PD. Computational consequences of temporally asymmetric learning rules: I Differential Hebbian learning. Journal of computational neuroscience. 1999;7(3):235–246.

60. Kingma DP, Ba J. Adam: A method for stochastic optimization. arXiv preprint arXiv:14126980. 2014;.

61. Dumoulin V, Visin F. A guide to convolution arithmetic for deep learning. arXiv preprint arXiv:160307285. 2016;.

62. Rolinek M, Martius G. L4: Practical loss-based stepsize adaptation for deep learning. arXiv preprint arXiv:180205074. 2018;.

63. Yamins DL, Hong H, Cadieu CF, Solomon EA, Seibert D, DiCarlo JJ. Performance-optimized hierarchical models predict neural responses in higher visual cortex. Proceedings of the National Academy of Sciences. 2014;111(23):8619–8624.

64. Nurminen L, Merlin S, Bijanzadeh M, Federer F, Angelucci A. Top-down feedback controls spatial summation and response amplitude in primate visual cortex. Nature communications. 2018;9(1):2281.

65. Candes EJ, Romberg JK, Tao T. Stable signal recovery from incomplete and inaccurate measurements. Communications on Pure and Applied Mathematics: A Journal Issued by the Courant Institute of Mathematical Sciences. 2006;59(8):1207–1223.

66. Shadlen MN, Newsome WT. The variable discharge of cortical neurons: implications for connectivity, computation, and information coding. Journal of neuroscience. 1998;18(10):3870–3896.

67. Van Vreeswijk C, Sompolinsky H. Chaos in neuronal networks with balanced excitatory and inhibitory activity. Science. 1996;274(5293):1724–1726.

68. Haider B, Duque A, Hasenstaub AR, McCormick DA. Neocortical network activity in vivo is generated through a dynamic balance of excitation and inhibition. Journal of Neuroscience. 2006;26(17):4535–4545.

69. Denève S, Alemi A, Bourdoukan R. The brain as an efficient and robust adaptive learner. Neuron. 2017;94(5):969–977.

70. Izhikevich EM. Which model to use for cortical spiking neurons? IEEE transactions on neural networks. 2004;15(5):1063–1070.

71. Friston K. The free-energy principle: a unified brain theory? Nature reviews neuroscience. 2010;11(2):127.

72. Bengio Y, Lee DH, Bornschein J, Mesnard T, Lin Z. Towards biologically plausible deep learning. arXiv preprint arXiv:150204156. 2015;.

73. Sze V, Chen YH, Yang TJ, Emer JS. Efficient processing of deep neural networks: A tutorial and survey. Proceedings of the IEEE. 2017;105(12):2295–2329.

74. Jouppi N, Young C, Patil N, Patterson D. Motivation for and evaluation of the first tensor processing unit. IEEE Micro. 2018;38(3):10–19.

75. Rotermund D, Pawelzik KR. Massively Parallel FPGA Hardware for Spike-By-Spike Networks. bioRxiv. 2018;doi:10.1101/500280.

76. Van Essen DC, Anderson CH, Felleman DJ. Information processing in the primate visual system: an integrated systems perspective. Science. 1992;255(5043):419–423.

77. Deneve S, Latham PE, Pouget A. Efficient computation and cue integration with noisy population codes. Nature neuroscience. 2001;4(8):826.

78. von Helmholtz H. Handbuch der Physiologischen Optik, Dritter Band; 1910.

79. Ghosh P, Sajjadi MSM, Vergari A, Black M, Schölkopf B. From Variational to Deterministic Autoencoders; 2019.

